# Linking genetic and environmental factors through marker effect networks to understand trait plasticity

**DOI:** 10.1101/2023.01.19.524532

**Authors:** Rafael Della Coletta, Sharon E. Liese, Samuel B. Fernandes, Mark A. Mikel, Martin O. Bohn, Alexander E. Lipka, Candice N. Hirsch

## Abstract

Understanding how plants adapt to specific environmental changes and identifying genetic markers associated with phenotypic plasticity can help breeders develop plant varieties adapted to a rapidly changing climate. Here, we propose the use of marker effect networks as a novel method to identify markers associated with environmental adaptability. These marker effect networks are built by adapting commonly used software for building gene co-expression networks with marker effects across growth environments as the input data into the networks. To demonstrate the utility of these networks, we built networks from the marker effects of ∼10,000 non-redundant markers from 400 maize hybrids across nine environments. We demonstrate that networks can be generated using this approach, and that the markers that are co-varying are rarely in linkage disequilibrium, thus representing higher biological relevance. Multiple covarying marker modules associated with different weather factors throughout the growing season were identified within the marker effect networks. Finally, a factorial test of analysis parameters demonstrated marker effect networks are relatively robust to these options, with high overlap in modules associated with the same weather factors across analysis parameters. This novel application of network analysis provides unique insights into phenotypic plasticity, and specific environmental factors that modulate the genome.

## Introduction

The global climate is changing at a faster pace than scientists initially predicted, and the impact on agriculture has been widely documented (Pörtner *et al*. 2022). Studies have shown that climate change has affected the yields and phenology of many crop species to different extents (Kukal and Irmak 2018; Ray *et al*. 2019) and increased the risk of global food insecurity (Springmann *et al*. 2016; Hasegawa *et al*. 2018). Therefore, plant breeders need to release varieties that are adapted to new climate conditions more quickly to help maintain a stable food supply in the future. One way to tackle this challenge is to exploit the remarkable phenotypic plasticity that plants have, i.e., the ability of one genotype to display different phenotypes under different growth conditions (Des Marais *et al*. 2013). A better understanding of the genetic basis of trait plasticity would allow plant breeders to make more informed decisions when implementing genomic selection programs.

Several different approaches have been used to understand phenotypic plasticity, each with its own advantages and disadvantages. Traditionally, breeders have used statistical approaches such as the Finlay-Wilkinson regression model (Finlay and Wilkinson 1963), AMMI models (Gollob 1968; Gauch 1988), and GGE biplots (Yan *et al*. 2000) to understand and visualize which genotypes have better performance and are more stable across different environments (Malosetti *et al*. 2013; Bernardo 2020). More recently, combining these approaches with genome-wide association studies (GWAS) allowed the identification of genomic regions associated with the overall stability of genotypes across different environments (Kusmec *et al*. 2017; Mangin *et al*. 2017; Gage *et al*. 2017). For example, GWAS with Finlay-Wilkinson regression slopes and deviations as explanatory variables showed that markers associated with stability are often found in regulatory regions of the maize genome (Gage *et al*. 2017). The main drawback of these approaches is that they take into account only the overall effects of environments on cultivar performance, without differentiating which specific environmental factor (e.g., temperature, precipitation, wind speed) is associated with a particular genetic marker. For this purpose, newer methodologies were recently developed. The Critical Environmental Regressor through Informed Search (CERIS) (Li *et al*. 2018, 2021a) algorithm identifies a combination of environmental factors and growth periods (e.g., photothermal time ∼50 days after planting) that is highly correlated with the mean performance of a trait (e.g., plant height) in different environments. Then, a Joint Genomic Regression Analysis (JGRA) is performed to estimate the effects of different genomic regions along this environmental gradient. In addition, Onogi *et al*. (2021) developed the Environmental Covariate Search affecting Genetic Correlations (ECGC), which identifies a similarity matrix based on environmental covariates that is highly correlated with a genetic correlation matrix between environments, and found that precipitation around sowing dates and hours of sunshine before maturity were both associated with variability in soybean yield.

Another common strategy to determine which genomic regions are associated with specific environmental factors is to conduct gene expression studies under controlled conditions as the analysis of gene expression profiles in plants reveal how certain genes increase or decrease their expression in response to different abiotic stresses (Wang *et al*. 2011; Jamil *et al*. 2011; Chawade *et al*. 2013). For example, through differential expression analysis, Frey *et al*. (2015) found over 600 heat-responsive and heat-tolerant genes across eight maize lines grown under mild and strong heat stress, and Fracasso *et al*. (2016) identified which genes of a sorghum genotype were associated with its increased susceptibility to drought. These data can also be used to generate networks composed of clusters of genes that have similar expression profiles across different samples or growing conditions, known as gene co-expression networks (Zhang and Horvath 2005). Individual clusters can then be correlated with variability of a trait of interest across the same conditions to identify relevant genes that are modulated across the samples and important for the trait of interest (Zhang and Horvath 2005; Langfelder and Horvath 2008). Gene co-expression networks benefit from increased statistical power to detect statistically significant associations compared to a gene-by-gene approach. For example, Li *et al*. (2021b) identified nearly 30 gene modules associated with variability of root traits grown under salinity stress, where one of them was associated with root length variability and contained approximately 3,000 genes with functions related to auxin transport, root development, and responses to abiotic stimuli. A major challenge with gene expression approaches is scaling them to breeding scales in field settings.

Here, we propose the concept of marker effect networks as a novel approach to identify regions of the genome associated with environmental adaptability. The idea is very similar to how gene co-expression networks work: genetic markers that have additive effect sizes for a phenotypic trait of interest varying similarly across different environments may be involved in the same biological process (in this case, phenotypic response to the different environments). This approach can leverage existing genetic data from breeding populations across multiple field environments (i.e., there is no need to design specific experiments for specific target environments), and provide a global understanding of how markers interact with each other when responding to different environments. Also, it can improve statistical power to detect associations with environmental factors (e.g., temperature, precipitation, wind speed) by correlating the eigenvalues of marker effects across environments for a group of markers with similar effects instead of individual markers.

In this proof-of-concept study, we calculated the effects of ∼10,000 markers on grain yield of 400 maize hybrids across nine environments and used the weighted gene co-expression network analysis (WGCNA) pipeline to build a marker effect network. We correlated network modules with the variability of specific weather parameters for the same set of environments and found markers associated with variation in grain yield due to evapotranspiration at the beginning of the growing season, precipitation in the middle of the season, and temperature towards the end of the growing season. We propose that marker effect networks can further our understanding of the genetic basis of trait plasticity and, thus, help breeders make informed decisions about which markers to use in their breeding programs based on the knowledge of specific environments.

## Materials and Methods

### Germplasm

Six maize inbred lines (B73, PHG39, PHG47, PH207, PHG35, and LH82) representing three heterotic groups were crossed among each other in a half diallel scheme. These 15 half diallel F_1_ crosses were self-pollinated to generate families of F_7_ recombinant inbred lines (RILs). A total of 333 RILs from 14 families were crossed among populations to generate 400 experimental maize hybrids (Table S1) that were subsequently phenotyped in nine environments.

### Growth environments and phenotypic data collection

The 400 maize hybrids were grown in two-row plots in a total of nine environments (i.e., Location × Year combination) across the U.S. Midwest in 2019 and 2020 with two replicates per environment (Table S2). These environments include five different locations (Bloomington, IL; two sites in Champaign, IL; St. Paul, MN; Janesville, WI) that were included in one or both years. Agronomic growth conditions for each environment are specified in Table S2. Plots were machine harvested. Plot grain weight and grain moisture at harvest were measured to determine grain yield (bu ac^-1^) normalized to 15.5% moisture. The number of hybrids for which yield data was collected ranged from 356 to 399 of the hybrids across the environments (File S1). Analysis of variance was conducted to assess significance of experimental factors using the following linear model with the R package lme4 (Bates *et al*. 2015):

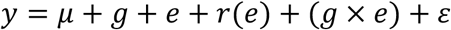

where *y* is grain yield values, *μ* is the overall mean, *g* is the fixed genotypic effect of a maize hybrid, *e* is the random environmental effect, *r*(*e*) is the random replicate effect within environment, *g* × *e* is the random effect of genotype-by-environment interactions, and *ε* is the residuals.

Best linear unbiased estimates (BLUEs) were calculated for the hybrids in each environment (File S2) using the following mixed linear model implemented in ASReml-R v4.1 (Butler *et al*. 2017):

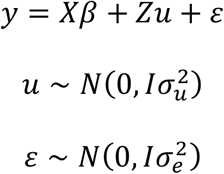

where *y* is the vector of grain yield data in one environment, *X* is the design matrix for the fixed genotype effects *β, Z* is the design matrix for the random replication effects *u, I* is an identity matrix, 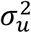 is the replication variance, and *ε* is the residual effect with variance 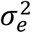 (Figure S1).

### Genotypic data

The six parents of this population and all of the RILs were genotyped with a custom Illumina Infinium 20k SNP chip (Files S3 and S4). To improve the quality of the genotypic data, we removed SNPs showing segregation distortion with the R package Rqtl v1.48 (Broman *et al*. 2003) using an FDR-adjusted p-value less than 0.05 from Chi-square test, and performed the sliding window correction described by (Huang *et al*. 2009) for each RIL family separately. The coordinates of each marker were converted to those of the B73 v4 reference assembly (Files S3 and S4). Genotypic data for each of the 400 maize hybrids were generated *in silico* by combining the two parental alleles at each locus. For cases when one of the parents had a heterozygous locus, the genotype of the respective marker in the hybrid was set to missing for both parents.

Given the structure of the population, it is expected that markers that are physically close to each other will be in high linkage disequilibrium (LD). To reduce the impact of LD on network clustering, the hybrid genotypic dataset was pruned by linkage disequilibrium (LD) (r^2^ > 0.9) within 100 kb windows with PLINK v1.9 (Chang *et al*. 2015). Additionally, markers with minor allele frequencies of less than 5% and missing data higher than 25% in the hybrid genotypes were removed, leaving a total of 10,334 high-quality markers for downstream analysis (File S5).

### Estimation of marker effects

Marker effects for grain yield in each environment were estimated in two ways. The first method used the RR-BLUP model implemented in the R package rrBLUP v4.6.1 (Endelman 2011):

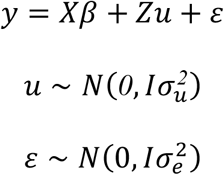

where *y* is the vector of hybrid BLUEs in a particular environment, *X* is the design matrix for the fixed effects *β, Z* is the design matrix for the random marker effects *u, I* is an identity matrix, 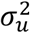 is the marker variance, and *ε* is the residual effect with variance 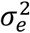 (Figure S2).

The second method used to estimate marker effects for grain yield was the unified mixed linear model (MLM) GWAS model implemented in GAPIT v3 (Yu *et al*. 2006; Wang and Zhang 2021):

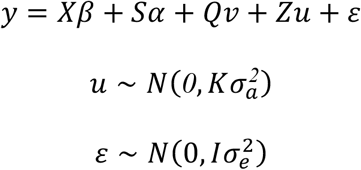

where *y* is the vector of hybrid BLUEs in a particular environment, *X* is the design matrix for the fixed effects *β, S* is the design matrix for the vector of fixed quantitative trait loci (QTL) effects *α, Q* is the matrix relating *y* to the fixed population effects *v, Z* is the design matrix for the random marker effects *u, K* is the kinship matrix, 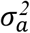 is the additive genetic variance, and *ε* is the residual effect with variance 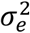 (Figure S3).

### Constructing marker effect networks

The marker effect networks were built with the R package WGCNA v1.70.3 (Zhang and Horvath 2005; Langfelder and Horvath 2008). This package was originally developed to construct weighted correlation networks from gene expression data across different samples. Here, instead of gene expression, we used the marker effects across different environments as our input data. Because marker effect data is inherently different from gene expression data with regard to scale and distribution, we tested a factorial combination of different parameters in building the networks and subsequently evaluated the impact on network quality (Tables S3 and S4).

The first set of parameters tested were different data normalization strategies to account for the differences in the range of effect values estimated in each environment, including minmax normalization (effects across environments were constrained between 0 and 1) and Z-score normalization (effects were centered on the mean and scaled by the standard deviation). In addition, we iterated through different coefficient of variation (CV) cutoffs in which markers with a CV lower than 0.05-0.25 after minmax normalization or 0.5-1.0 after Z-score normalization were removed to improve the scale-free topology model fit (signed R^2^; Zhang and Horvath 2005) of the network.

Marker effect networks were then built with default values from the WGCNA pipeline using a soft thresholding power of 24, except for two parameters of the *cutreeDynamic* function responsible for defining the modules of the network. The first parameter was *minClusterSize*, which determines the minimum number of markers to be present in a module. Networks were created with this parameter set to 25, 50, or 100 markers. The second parameter tested was the *pamStage* that was set to either ‘on’ or ‘off’. If on, a Partitioning Around Medoids (PAM)-like step is used to assign outlying markers to the nearest modules. Disabling this option may result in “cleaner” modules, but the number of markers not assigned to any module may increase (Langfelder *et al*. 2008).

### Assessing marker effect network connectivity

The quality of each network was assessed based on the overall connectivity and cohesiveness of their modules based on three metrics: clustering coefficient (i.e., tendency of a marker to associate with its own module), kDiff (i.e., number of connections of a marker with other markers within its own module minus the number of connections to markers outside its module), and kRatio (i.e., ratio between kDiff and total number of connections). An ideal network would have all modules with a high clustering coefficient, positive kDiff, and kRatio close to 1, meaning that markers within each module are more connected with each other than with markers in other modules.

### Assessing linkage disequilibrium within modules

Markers were pruned for LD within a 100 kb window prior to network construction. However, given the population structure of the 400 hybrids, LD between markers outside of the 100kb window or on different chromosomes can occur. To assess LD within modules, PLINK v1.9 (Chang *et al*. 2015) was used to calculate r^2^ between each pair of markers in a module using the hybrid genotypic data, regardless of the chromosomes on which they were located.

### Obtaining weather factors and indices

The R package EnvRtype v1 (Costa-Neto *et al*. 2021) was used to extract 17 weather factors (Table S5) across the nine environments evaluated in this study (Table S2) in 3-day intervals from planting date to the end of the season. The last time interval was set to 151 days after planting to have each environment with an equal number of intervals, resulting in a total of 867 environmental indices (File S6). Pearson correlation between all environmental indices was calculated using the cor function in R. Principal component analysis (PCA) was performed in R using the prcomp function to reduce the dimensionality of this highly correlated set of weather indices.

### Correlation between marker effect network modules and weather indices

To understand how different groups of markers relate to weather factors, Pearson correlations were calculated between the eigenvalues for each module of a marker effect network and a principal component of the environmental indices described above out to the ninth principal components based on visualization by a scree plot (Figure S4). P-values were adjusted with a false discovery rate (FDR; Benjamini and Hochberg 1995) correction to account for multiple testing. The marker composition of modules from different networks that were significantly associated with the same principal component were visualized with upset plots from the R package UpSetR v1.4 (Conway *et al*. 2017).

### Data and code availability

The datasets analyzed in this study are available as Files S1-S6 in the Data Repository for the University of Minnesota (DRUM; https://conservancy.umn.edu/drum). All scripts necessary for data analysis are available on GitHub at: https://github.com/HirschLabUMN/meffs_networks.

## Results and Discussion

### Evaluating input data of networks

The main goal of marker effect networks is to identify genetic factors that contribute to environmental responsiveness in a coordinated way. Specifically, the analysis aims to identify markers with correlated effect sizes on a particular trait across environments that are modulated by specific environmental factors. To develop these networks, we need phenotypic data on a set of genotypes grown over a diverse range of environmental conditions from which marker effects are estimated. To this end, we grew a population of 400 maize hybrids in nine environments and collected grain yield data. The population structure of these lines resembles the type of structure breeders commonly use in their programs making it an ideal set of germplasm on which to test the utility of developing marker effect networks and testing correlations with environmental factors. Significant effects of genotype, environment, and genotype-by-environment factors were all observed in the analysis of variance (ANOVA), all of which are necessary sources of variation to effectively build marker effect networks (Table S6).

The input into the networks is the additive effect of each marker in each environment for each genotype. There are many models available to estimate these effects. We tested the RR-BLUP and GWAS models due to their relative simplicity and broad use by breeding programs. The estimated effects across all environments were highly correlated between the two models (r = 0.92), and the genomic estimated breeding values (GEBVs) were also highly correlated with the best linear unbiased estimates (BLUEs) of the hybrids (r = 0.85 for both models; Figure 1a). Given the highly correlated results across these standard methods for estimating marker effects, substantial differences in the networks constructed from either set of marker effects were not anticipated, and as shown below, this is the case.

**Figure 1.**
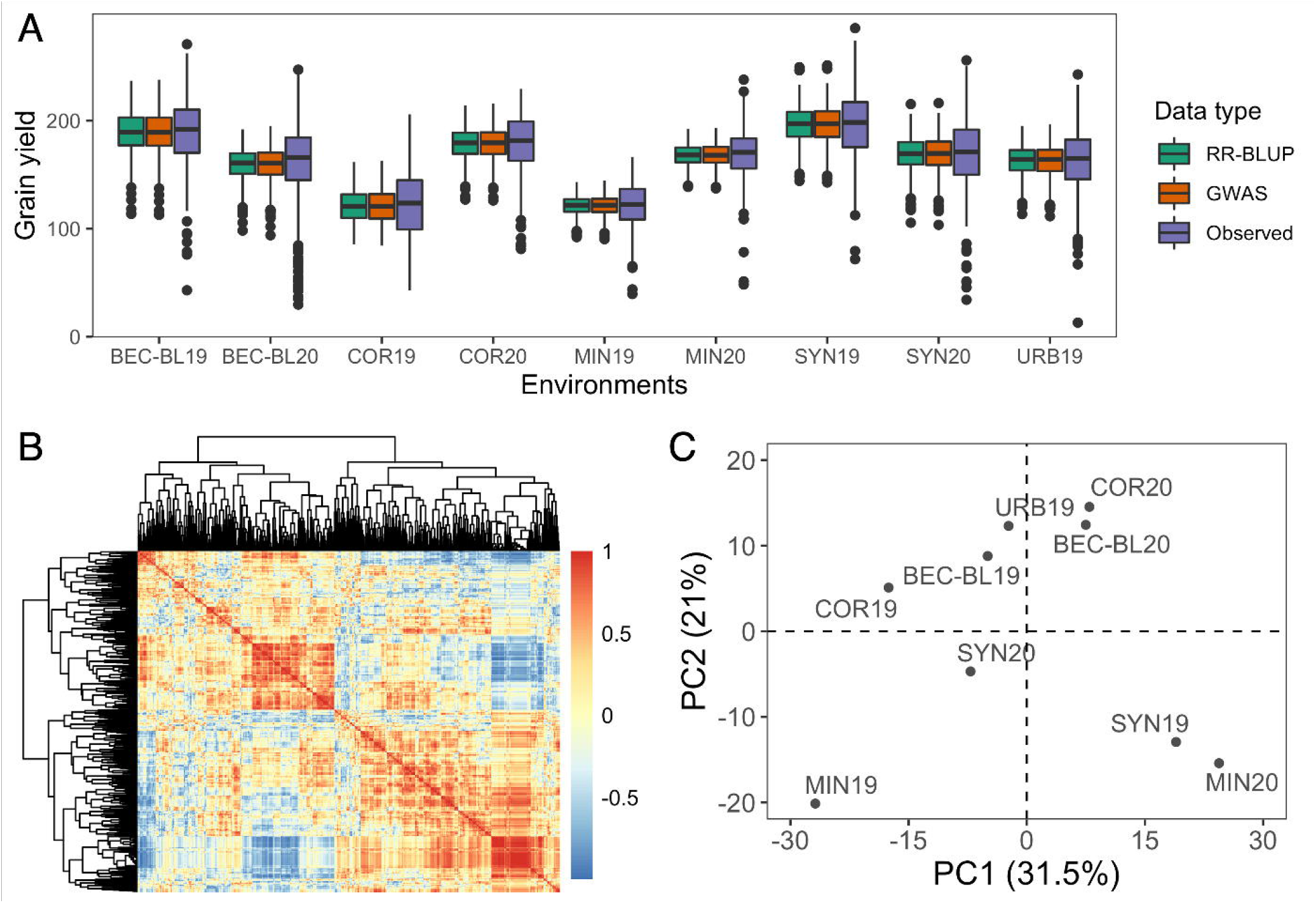
Experimental variation in the input data into the marker effect networks. (a) Distributions of genomic estimated breeding values (GEBVs) from two different models and the observed phenotypic values across nine environments. (b) Heatmap of correlations among 867 environmental factors from 51 time intervals throughout the growing season across nine environments. (c) Principal component analysis of the 867 environmental factors across the nine growth environments. Each data point in the plot represents a single environment labeled by its environmental code (Table S2).

An important factor to consider when building marker effect networks is the number and diversity of the environments. This idea is similar to gene co-expression networks, where many samples with sufficient variation are needed to identify real signals of co-expression (Li *et al*. 2015). Usually, in network analysis, the more samples (or in our case environments) that are available, the higher are the chances of finding biologically meaningful associations between network modules and traits (or environmental factors) of interest (Li *et al*. 2015). We evaluated the diversity of our nine growth environments by analyzing 17 environmental factors across 51 time intervals spanning the entire growing season. The 867 environmental factors were highly correlated (Figure 1b), and as such, we reduced the dimensionality of the data by conducting principal component analysis. The first and second principal components provided a clear separation of the growth environments (Figure 1c). Thus, it is likely that at least a portion of the significant environment and genotype-by-environment interaction factors in the ANOVA were driven by weather factors and not just soil types or agronomic management differences across environments, and these weather differences could potentially be correlated with variability in marker effects across the environments.

### Building marker effect networks

There are many different factors that can be modified and optimized in conducting network analyses (Rao and Dixon 2019). Optimized parameter combinations have been previously assessed for building gene co-expression networks (Langfelder and Horvath 2008). With marker effect networks we are using a novel data input type for the network analysis, and therefore tested different marker effect estimation methods (GWAS vs. RR-BLUP), data normalization strategies (minmax and Z-score), a range of coefficient of variation filters, and soft thresholding power that all could potentially affect the scale-free topology fit.

In total, we assessed the impact of 36 different parameter combinations on the scale-free topology fit (Table S3). With many of these options, there is a balance between the power needed to maximize the scale-free topology fit and the number of markers that are retained and ultimately assigned to meaningful modules. For example, higher CV cut-offs usually led to smaller power needed to reach a good scale-free topology fit (Figure 2a), but also considerably decreased the total number of markers remaining to build the network (Figure 2b). For many of the parameters, there was a minimal observed difference with regard to the minimum power needed to reach a good scale-free topology fit. Though marker effects estimated via RR-BLUP resulted in better signed R^2^ values (Zhang and Horvath 2005) using slightly smaller power than GWAS, and similar results were observed when comparing minmax and Z-score normalization strategies (Figure 2a). Across all the parameter options, we obtained good model fit (signed R^2^ > 0.7) only when using very high powers (around 20 for most cases; Figure 2a), which is considerably higher than powers typically used to build gene co-expression networks (Zhang and Horvath 2005; Amrine *et al*. 2015; El-Sharkawy *et al*. 2015; Tan *et al*. 2017; Sharma *et al*. 2018). This result reflects the differences in scale and distribution of input values between marker effect and gene expression data. The optimal combination of parameters with regard to scale-free topology fit for our marker effect data was to use a Z-score normalization with a CV cut-off of 0.05 and a power of 24. This balanced a good scale-free topology fit with the total number of markers available to build a network.

**Figure 2.**
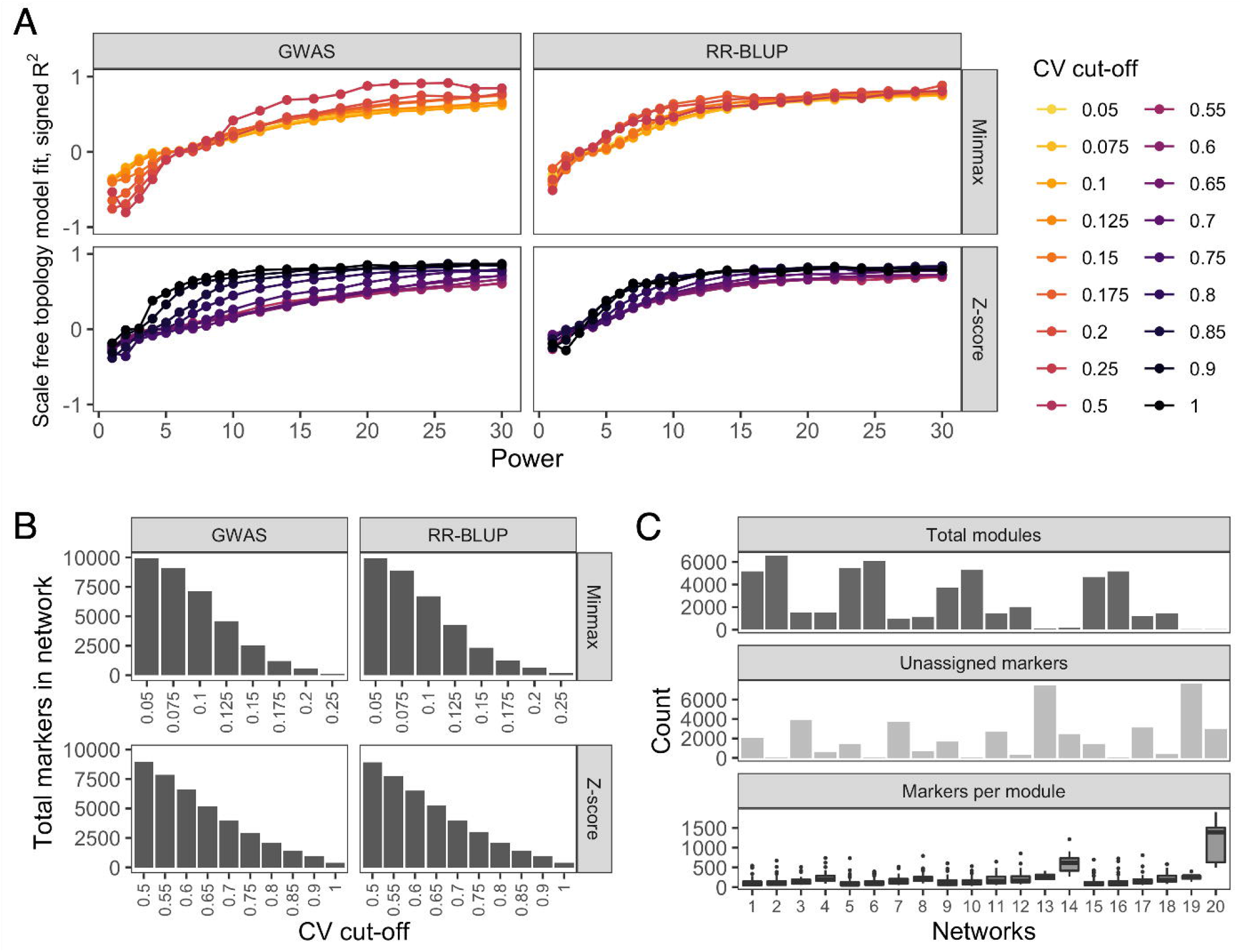
Characteristics of module defined marker effect networks built with different parameter combinations. (a) Scale free topology fit (signed R^2^; (Zhang and Horvath 2005)) for different powers. Each panel represents a different combination of marker effect estimate and normalization methods, while the line colors represent different coefficient of variation (CV) thresholds. (b) Total number of markers remaining in a network after applying different CV thresholds. (c) Number of modules, unassigned markers, and markers assigned per module for different networks. Parameter combinations for networks 1-20 can be found in Table S4.

After building the networks, the modules within those networks need to be defined. As with building the networks, there are a number of options that can be iterated when defining the modules within the network. To find the optimal combination we tested the minimum number of markers allowed in a module (25, 50, or 100) and whether or not to assign outlying markers to the nearest modules (PamStage on or off, respectively) in combination with different methods for estimating marker effects and data normalization strategies as there may be an interaction between parameter options within these different stages of the analysis. In total, we aimed to generate 24 different sets of network defined modules of co-varying markers from a factorial combination of all network parameters described above. However, we were not able to define a single module for four networks when using a minimum of 100 markers per module and ended up with a total of 20 different module defined networks (Table S4). The total number of modules per network decreased considerably as the minimum number of markers required to be in a module increased (Figure 2c). For example, 61 to 81 modules per network were defined with a minimum of 25 markers (median markers per module ranging from 64.5 to 103), compared to only 6 to 13 when at least 100 markers were required in each module (median markers per module ranging from 276.5 to 1450). Disabling the pamStage option resulted in a larger number of markers not assigned to any module, as was expected (Figure 2c). As with the power assessment above, the method used to estimate marker effects and type of data normalization did not show any major impact on the definition of modules with regards to the number of modules or average module size.

### Marker effect networks have lower overall connectivity

In addition to assessing the number of modules and markers per module we also assessed connectivity metrics such as clustering coefficient, kDiff, and kRatio as additional metrics of network quality across the 20 module defined networks described above. Most modules within each network had markers with clustering coefficients closer to zero than to one Figure 3a, indicating that their tendency to associate with other markers in the same module is relatively weak. Similar results were obtained when evaluating the kDiff metric Figure 3a. In this case, the kDiff distribution is centered around zero, meaning that, on average, markers have a similar number of connections within and outside their modules. Across the metrics for assessing within module connectivity, lower connectivity was observed for marker effect networks compared to what is typically seen in co-expression networks (Figure 3a; Mähler *et al*. 2017; Hoopes *et al*. 2019; Sancho *et al*. 2022). One possible explanation may be the high power used to fit scale-free topology, which lowers the mean and median connectivity of the modules (Figure S5). The impact of network parameters on the quality of a network was more evident when kDiff was scaled with the total number of connections of each marker (i.e. kRatio). Increasing the minimum number of markers per module and disabling the pamStage resulted in slightly better kRatio (i.e., a higher proportion of markers connected to markers in their own modules).

**Figure 3.**
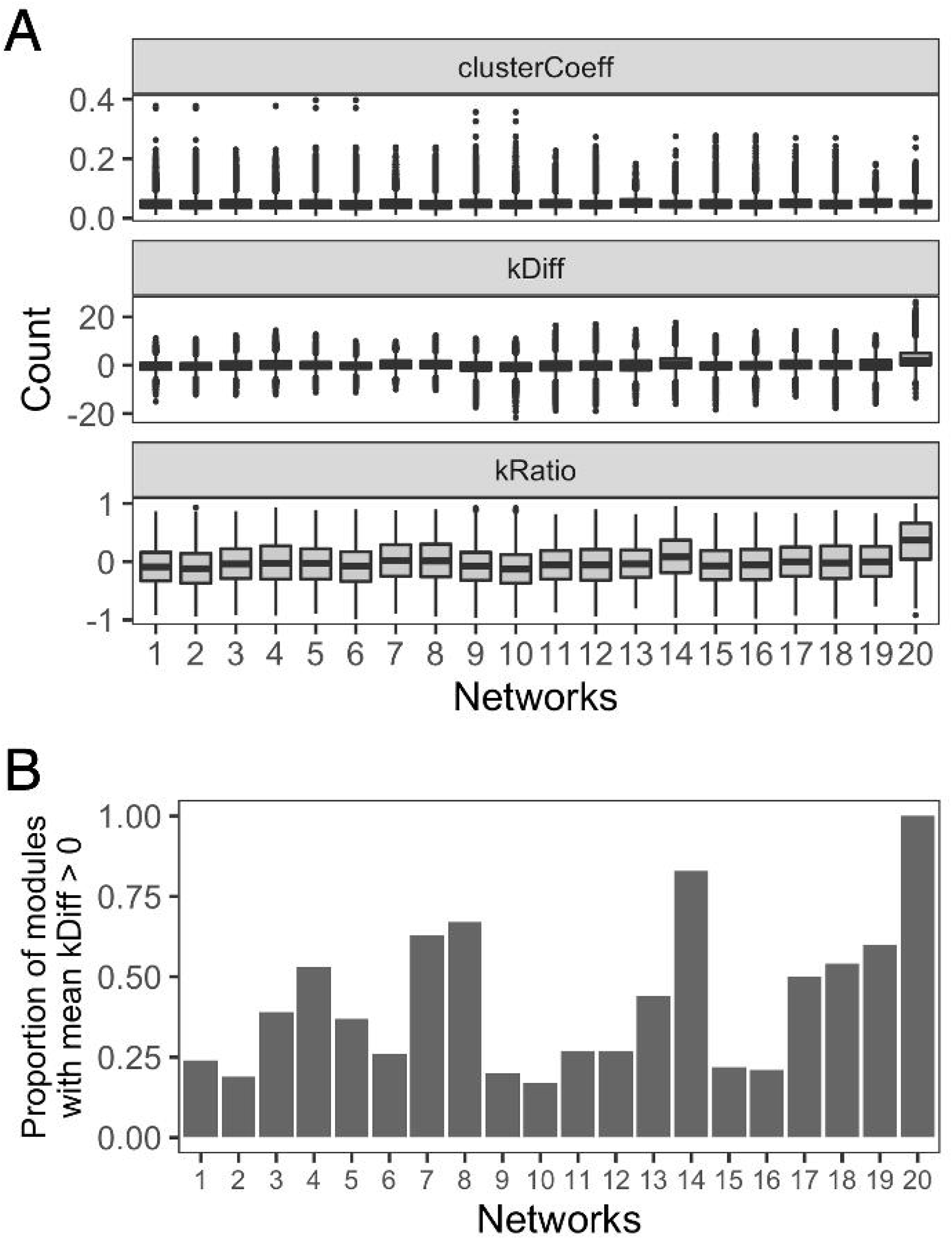
Quality assessment of marker effect networks built with different parameter combinations. (a) Distribution of values for different network connectivity metrics (clusterCoeff, kDiff, and kRatio). (b) Proportion of modules with average kDiff > 0 within each network. Parameter combinations for networks 1-20 can be found in Table S4.

These results do not imply that the entire network is noisy. Indeed, there is a range of intramodular connectivity scores within each of these networks, and those modules with higher intramodular connectivity (Figure 3b) will likely have more biologically meaningful connections with which specific environmental factors can be associated. Additional optimization of the WGCNA pipeline may further improve network connectivity and reduce the overall noise of network connectivity. Furthermore, there are alternative network building software that use different algorithms for building networks and defining modules (e.g., Camoco (Schaefer *et al*. 2018) and machine learning approaches (Li *et al*. 2015)) that may be able to achieve higher overall connectivity and improve the ability to find biologically meaningful signals within individual modules.

In gene co-expression networks, gene ontology analysis can be done to evaluate whether there is an enrichment of genes with the same function clustered in the same module (Langfelder and Horvath 2008). Similar validation is not possible in marker effect networks because of LD among markers and not knowing which marker is the causative variant. One possibility would be to overlap known QTLs regions to markers found by the network modules. This approach has been applied with gene co-expression network analysis (Schaefer *et al*. 2017, 2018; Li *et al*. 2021b), but will be highly dependent on the amount of information available for that population of individuals and the trait of interest. Another possibility is to run genomic prediction models with and without those markers on the same environments and see which set has better accuracy, but further studies would be necessary to determine the utility of this validation approach.

### Marker effect networks are not driven by genetic linkage disequilibrium

One key concern when building marker effect networks was that markers within a module would be highly correlated simply due to linkage disequilibrium (LD), especially for populations commonly used in breeding programs where the linkage blocks are relatively large. To minimize this possibility, we first filtered out markers that were in high LD within 100 kb. However, markers can still be in the LD with each other without being in physical proximity. To assess if this was the case, and if markers in genetic linkage were driving the connectivity of modules within networks, we calculated LD among all of the markers assigned to each of the modules regardless of chromosome number or position. Taking a closer look at one module (called firebrick2 by WGCNA) of Network 1 (Table S4), 90.1% of all marker connections within that module did not have an LD value (r^2^) above 0.9 (Figure 4a). Analyzing one module at a time allowed us to visualize the small portion of the intramodular connectivity driven by LD (Figure 4b). Similarly, low rates of LD within modules were observed across all modules in network 1, and across all of the 20 module-defined networks, with an average of only 3.2% and 2%, respectively, of marker connections being established due to LD (Figure 4c). This result suggests that the majority of the co-varying effects across environments were due to regions of the genome that are part of the same biological response processes rather than simply being a product of being genetically linked with biologically relevant regions of the genome.

**Figure 4.**
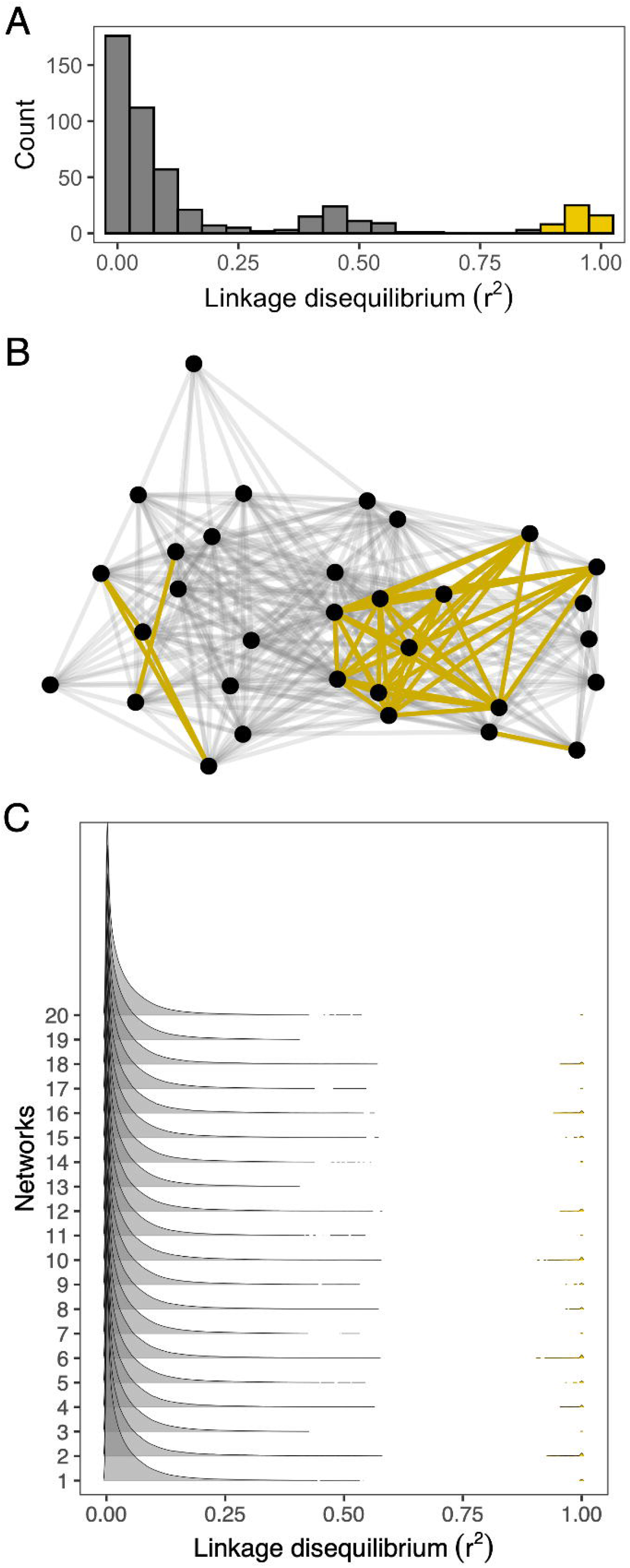
Linkage disequilibrium (LD) between markers within a module (a) LD distribution of markers in module firebrick2 of network 1 (Table S4) with the amount of marker connections in high LD (r^2^ > 0.9) highlighted in yellow. (b) Intramodular connectivity of the firebrick2 module where circles represent markers, yellow lines connect markers in high LD (r^2^ > 0.9), and gray lines connect markers not in high LD. (c) LD distribution of markers within modules across all 20 networks. Marker connections in high LD (r^2^ > 0.9) are highlighted in yellow.

### Marker effect networks identify environmental factors associated marker effect variation across environments

We next tested if the variation of marker effects in each module (represented by its eigenvalues) can be associated with variation in specific environmental factors, such as temperature or precipitation. This is conceptually similar to correlating gene expression patterns of a module to certain characteristics of the samples used to create the network, such as variations in plant phenotypes due to stress (Guo *et al*. 2020; Li *et al*. 2021b). An important network quality factor to consider before making these associations is to determine whether the marker effect patterns are driven by the presence of one or a few extreme environments, as this can impact the scope of environments in which these associations are relevant. To assess this within Network 1 (Table S4), we plotted the eigenvalues for all of the modules within the network (Figure 5), and observed a lot of variation of module eigenvalues across environments and no patterns that implicate specific environments driving assignment of markers into any of the module. Similar results were observed across the other 19 module-defined networks (Figure S6).

**Figure 5.**
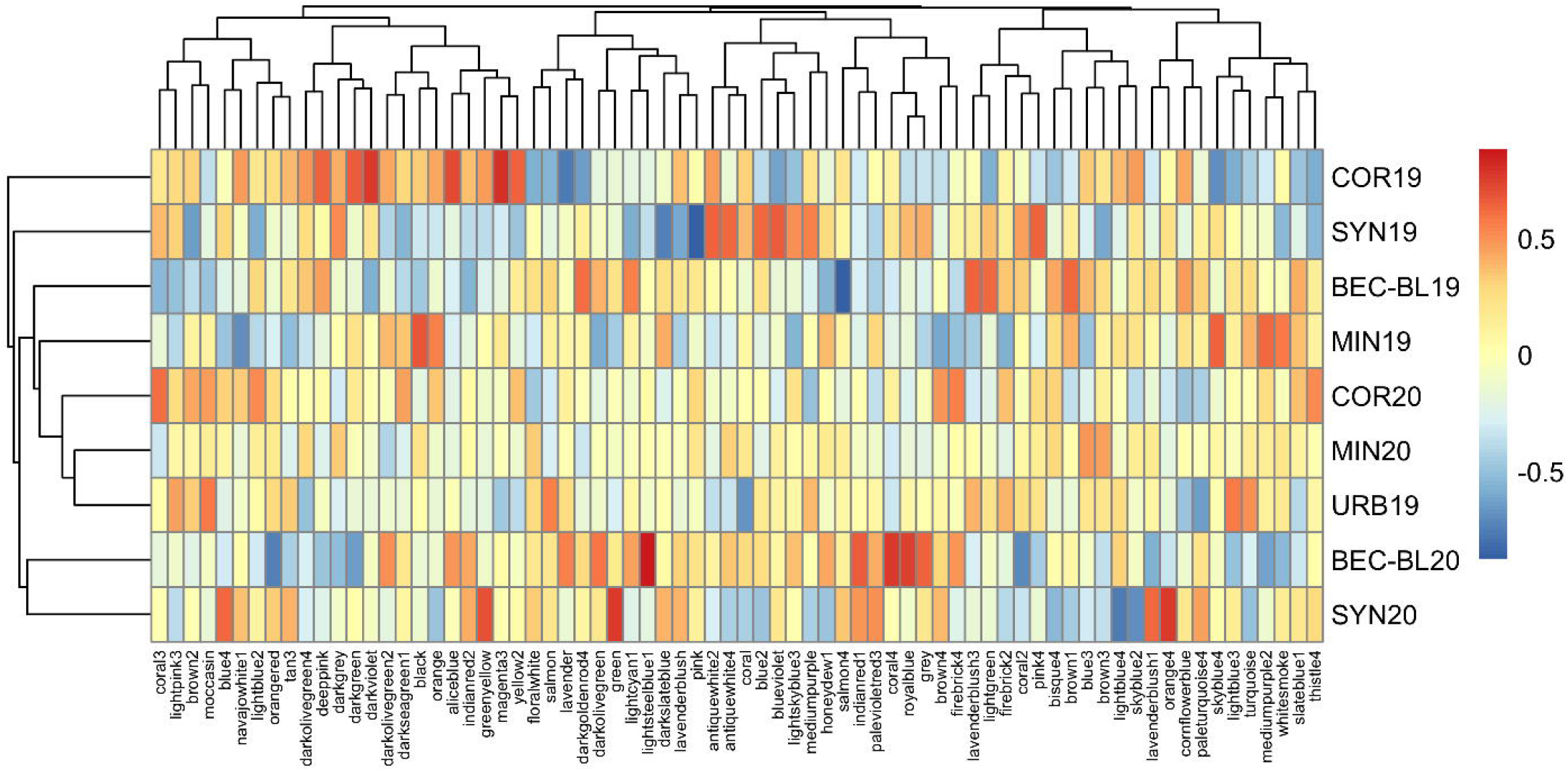
Heatmap of module eigenvalues (columns) from Network 1 across environments (rows). Higher eigenvalues are represented in red, while lower eigenvalues are in blue.

To make the association between modules and environmental factors, we obtained data for 17 weather factors in 3-day intervals across the entire growing season for a total of 867 individual weather factors per environment. Due to the highly correlated nature of many of these factors (Figure 1b), we performed a PCA to reduce the dimensionality of this data (Figure 1c). The first principal component (PC1) explained 31.4% of the variation in the 867 weather factors, while the last principal component (PC9) that we evaluated explained less than 0.01% of the variation in weather data represented by the original 867 weather factors (Figure S4). We correlated each principal component (PC) across the growing environments with the eigenvalues of each marker module. For simplicity, hereafter we are going to describe results from only Network 1 (Figure 6a; Table S4), and results for all other networks are visualized in Figure S7. After correcting for multiple testing, we observed that three modules from Network 1 were highly correlated with three distinct PCs (module firebrick2 and PC9: r = 0.91, p-value = 0.05; module lavenderblush3 and PC3: r = -0.89, p-value = 0.09; module brown3 and PC4: r = -0.89, p-value = 0.1). Such near statistically significant p-values are explained by the relatively high amount of noise present in the network, as indicated by the connectivity metrics, and the large number of tests (n = 648) done to associate every module with all nine PCs, which places a higher degree of multiple testing correction onto these tests.

**Figure 6.**
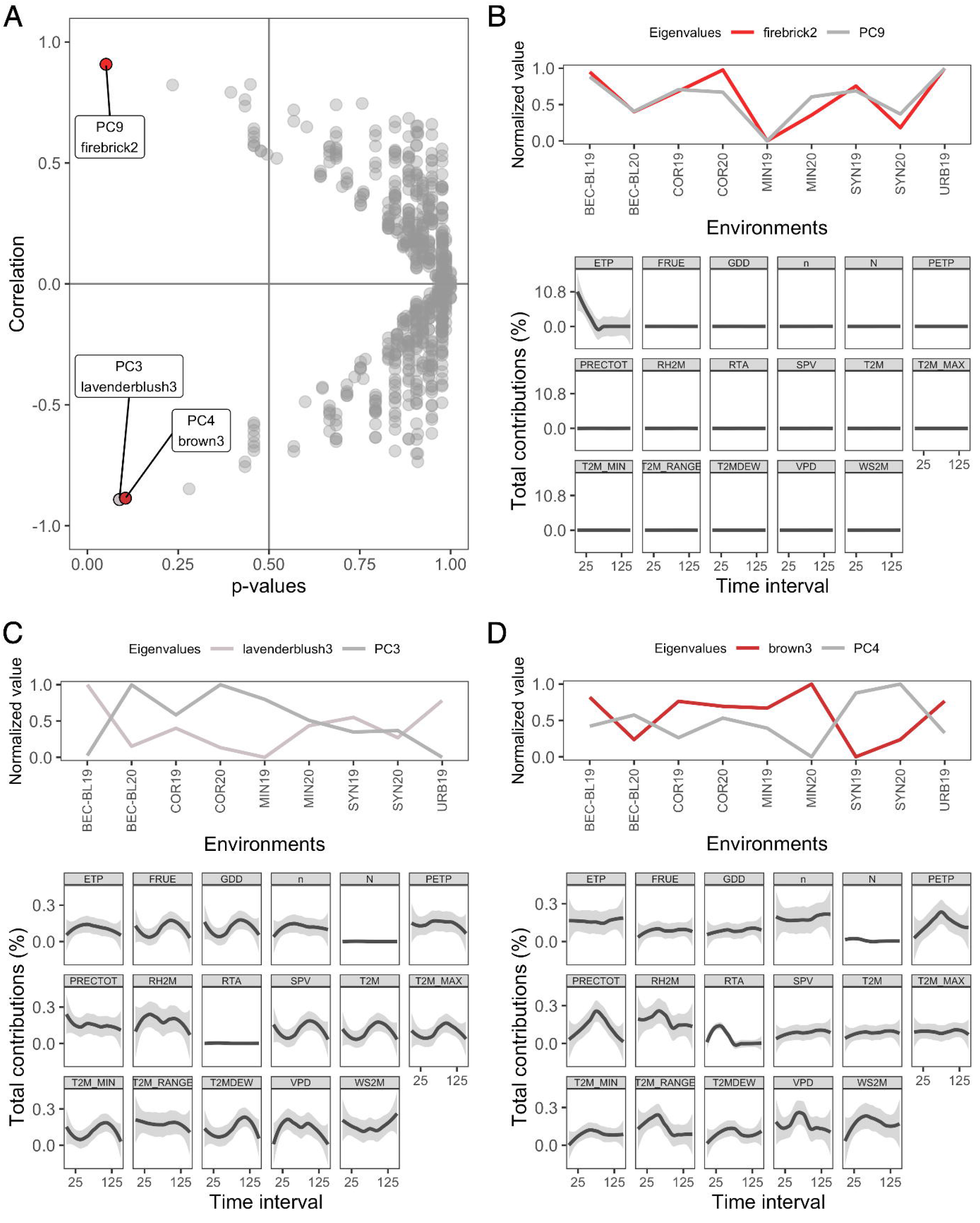
Correlation of marker effect network with eigenvalues and principal components (PC) of environmental factors across environments. (a) Correlation coefficient (y-axis) and significance (x-axis) between the eigenvalues of each network module and PC (represented by circles) with the three highest correlations highlighted. (b-d) Module and PC eigenvalues across growth environments (upper panel) of the three highest correlations and loess curves representing the contributions of individual environmental factors to the loadings of the respective PC (lower panel).

To understand which environmental factors were driving the variation of these significantly correlated PCs we investigated the loadings of each of the PCs. Interestingly, we found that PC9 is exclusively driven by variability in potential evapotranspiration (mm/day) in the beginning of the growing season (Figure 6b). Thus, the 32 markers in the firebrick2 module are candidates for capturing variation in grain yield due to changes in evapotranspiration rates across environments. For PC3 and PC4 the loadings were not as specific to a single weather factor at a single window of time. Rather, markers associated with PC3 tend to capture more grain yield variability due to differences in temperature (T2M, T2M_MAX, T2M_MIN, GDD) towards the end of the season (Figure 6c), and markers associated with PC4 tend to capture more grain yield variability due to differences in precipitation (PETP, PRECTOT, RH2M) in the middle of the growing season (Figure 6d). These environmental correlations provide valuable information in identifying regions of the genome that co-vary in effect sizes in response to weather patterns in the growing environment at different points in the growing season.

### Marker effect networks are relatively robust to changes in network parameters

In this study, we generated 20 different networks based on a combination of different network construction parameters. Since network structure and connectivity metrics varied between these networks (Figures 2 and 3), we also wanted to test the impact these parameter choices had on the biological connections that were observed. For this purpose, we compared the markers contained in modules across the different networks that were associated with the same environmental PC. Across the 20 networks, we observed 12 near significant correlations (p-values <= 0.1) between a module and an environmental PC. These significant correlations involved five different PCs, with four of the PCs having correlation to multiple modules across different networks (Table S7). The fact that changing parameters identified new associations with PCs suggests that some parameter combinations are more ideal to cluster markers associated with that PC than others. For example, in networks where modules are forced to be large, the signal that could have been captured by a smaller module is diluted. This is corroborated by the fact that 6 out of 8 eight networks with significant associations required only 25 markers per module, and no significant correlations were observed for networks with a minimum of 100 markers per module. Thus, while parameters that forced larger modules had generally higher connectivity scores, the biologically meaningful connections with regards to modulation by specific weather factors are lost as these markers are unable to be assigned to a module that meets the required minimum module size.

For the modules associated with the same PC, there is a high overlap between markers contained within the modules. For example, among the 56 unique markers from modules representing four different networks associated with PC9, 19 (34%) were shared, or in high LD to a shared marker, across all networks (Figure 7a). High overlap of markers was also observed for those modules associated with PC4 (Figure 7b; 52% overlap) and PC7 (Figure 7c; 94 % respectively). In contrast, we did not observe high overlap of the markers associated with PC5 across networks 3 and 4 (Figure 7d; 1.5% overlap). This may be a result of the much higher number of markers per module in those networks which may be noise in the module or biologically meaningful connections that were unable to be captured under specific parameter choices. Taken together, these results suggest that marker effect networks are relatively robust to changes in network parameters and that these overlapping markers may play a more important role in the module association with a PC.

**Figure 7.**
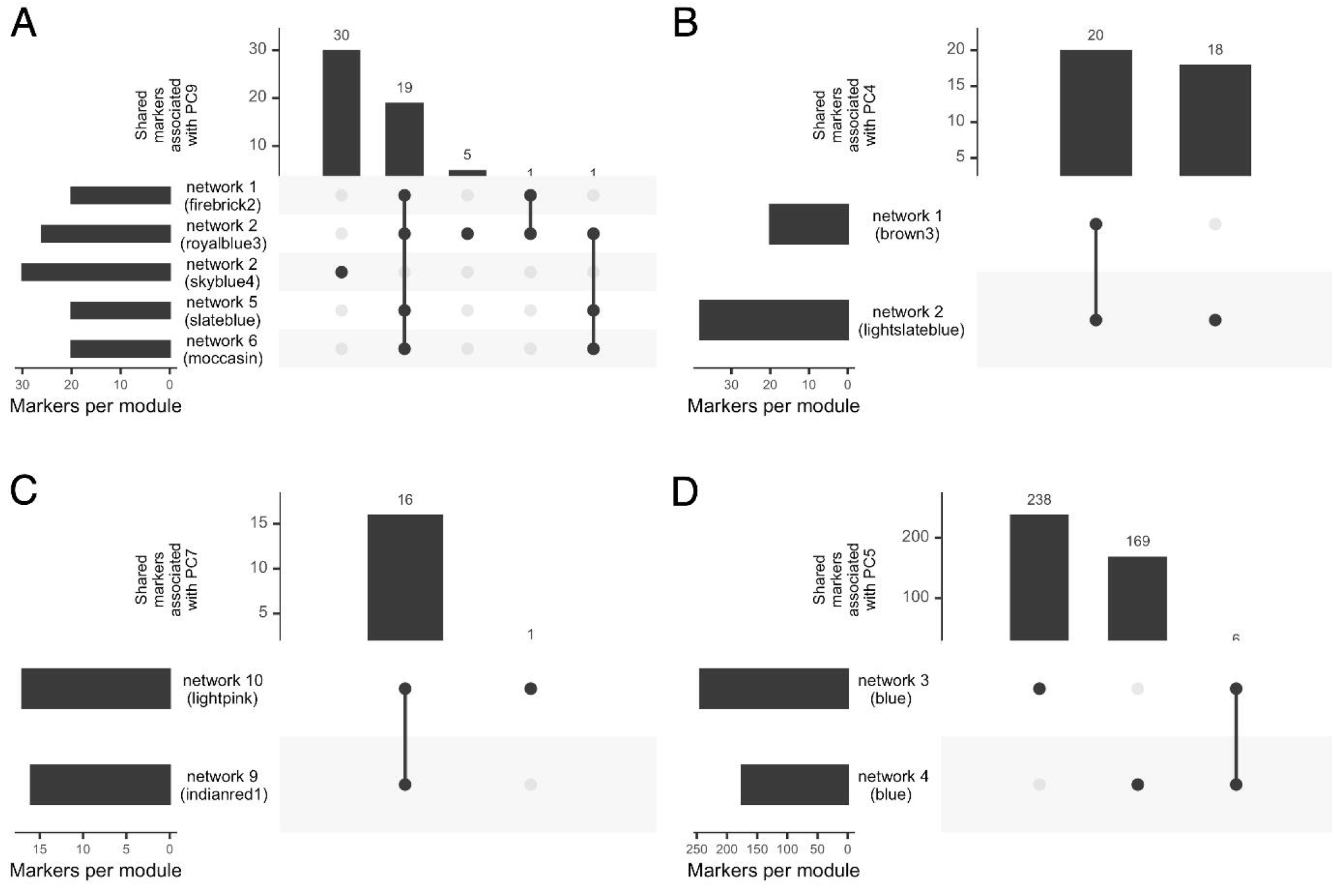
Upset plots with the number of overlapping markers from different modules correlated with the same principal component (PC). Markers across modules with linkage disequilibrium values greater than r^2^=0.9 were counted as overlapping between the modules. Environmental factor principal components with significant correlation to modules in more than one network included PC9 (a), PC4 (b), PC7 (c), and PC5 (d).

## Conclusions

As agricultural production continues to face changing and increasingly challenging growth conditions, understanding the factors that drive phenotypic plasticity in response to environmental factors will be important. Here we document a novel application of network analysis, in which the effects of markers across environments are used to build networks that allow for the identification of markers in the genome that co-vary in response to the environment. The outputs of these networks provide breeders with an atlas of markers that are most useful to use in genomic selection programs in different environmental contexts (Figure 8). If a breeder knows a group of markers captures a lot of variation across environments due to a specific environmental factor, they may choose to add or remove these markers based on how much their target environments vary for that environmental factor. For example, if a breeder is making predictions only across irrigated environments, they may choose to remove markers associated with precipitation and allocate resources to those markers associated with non-precipitation related weather variables. These associated modules in networks also provide valuable candidate genes for subsequent physiological studies.

**Figure 8.**
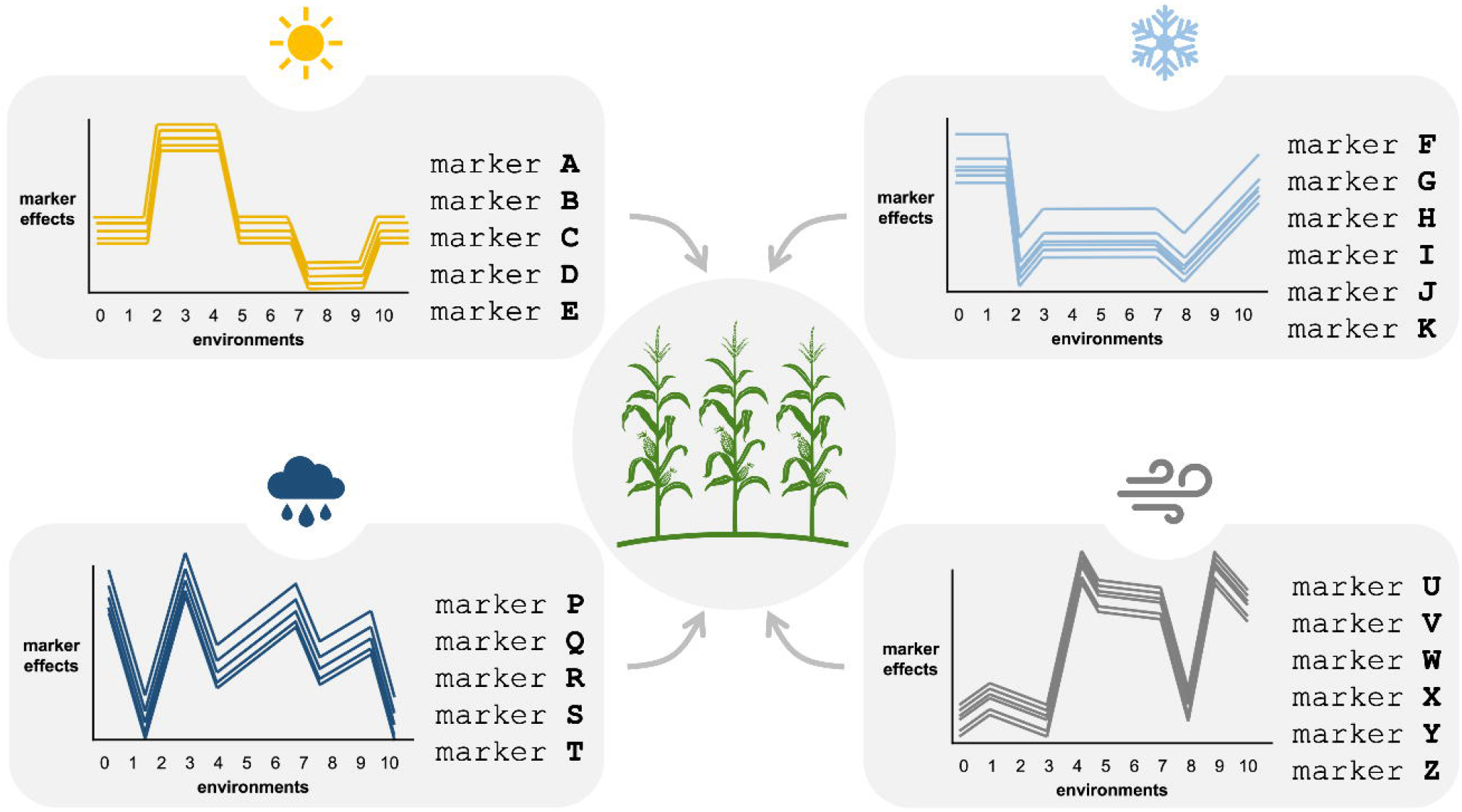
Conceptual maker effect network atlas. Genetic markers that have their effects covarying across different environments are grouped together because their response to these environments may be modulated by the same environmental factor, such as amount of sunlight, precipitation, temperature, and wind speed. Knowledge of how genetic and environmental factors interact would allow breeders to better understand the plasticity of their traits of interest, and fine tune their genomic selection programs. Corn image by Annette Spithoven from thenounproject.com.

Overall, our study demonstrates that marker effect networks are a promising new way to understand the genetic basis of trait plasticity. These networks are built from data that is commonly generated in breeding programs and so are readily available across a range of species and growing environments. The networks provide high resolution understanding of the environmental factors that are driving trait plasticity across complex environments. The results of these networks provide important lists of markers for future in-depth physiological studies (e.g., understanding mechanisms of water deficit response) and for direct use in breeding applications through inclusion in genomic prediction models.

## Acknowledgments

We thank Bayer Crop Science, Corteva Agriscience, Syngenta, and Beck’s Hybrid Corn Seed for providing in-kind support through growing locations for the multi-environment yield trial data used in this study and DOW AgroScience (now Corteva Agriscience) for providing in-kind support through the custom Illumina Infinium 20k SNP chip. We thank the Minnesota Supercomputing Institute at the University of Minnesota (http://www.msi.umn.edu) for providing resources that contributed to the research results reported in this article.

