## Supplemental Figures 1-7 for "Linking genetic and environmental factors through marker effect networks to understand trait plasticity"

**BEC-BL19**

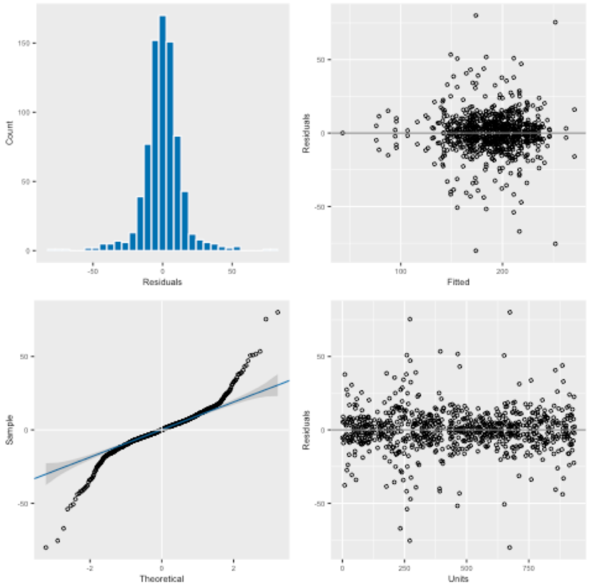


**BEC-BL20**

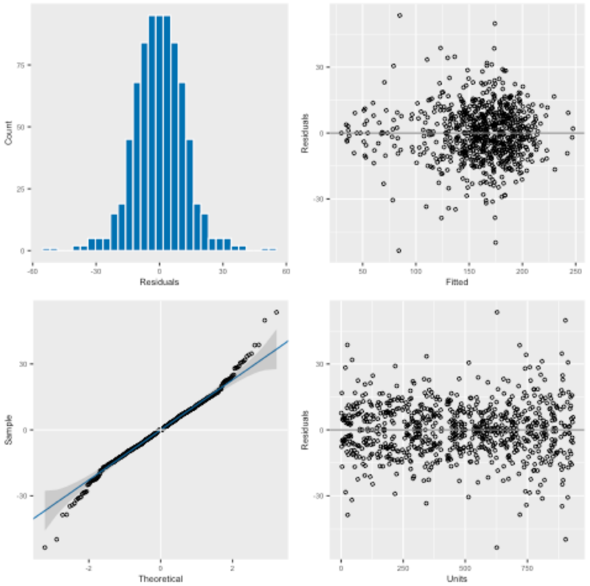


**COR19**

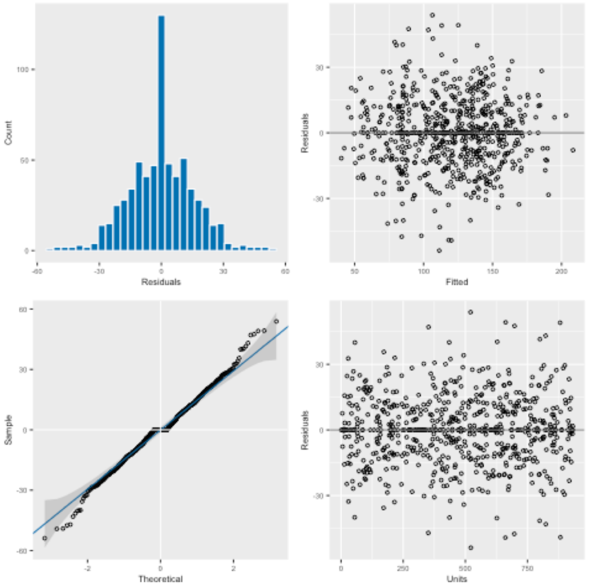


**COR20**

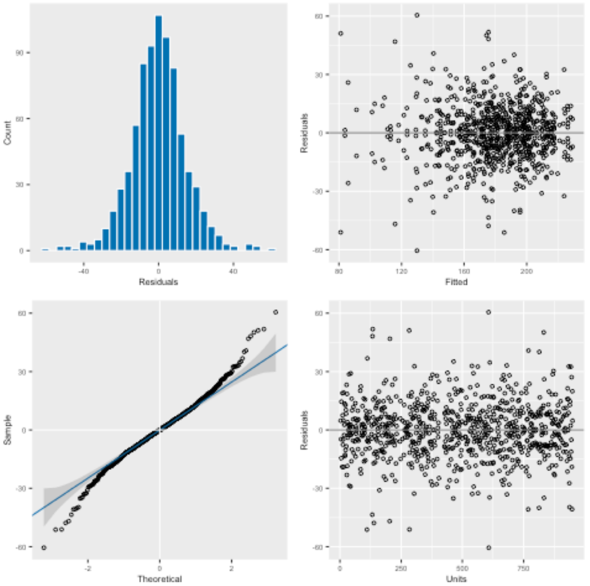


**MIN19**

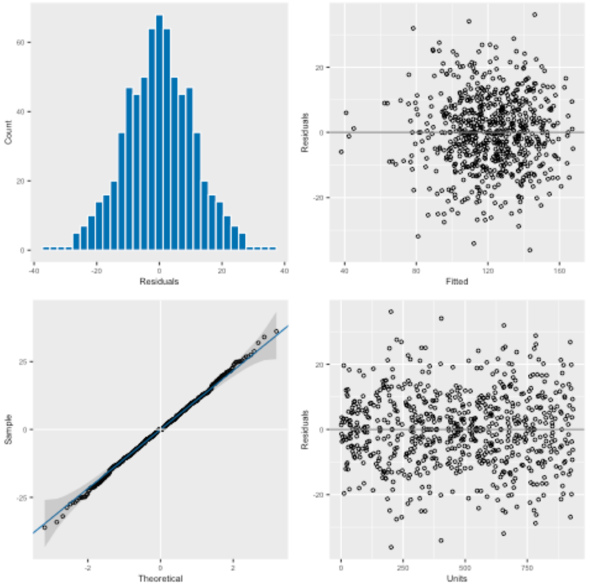


**MIN20**

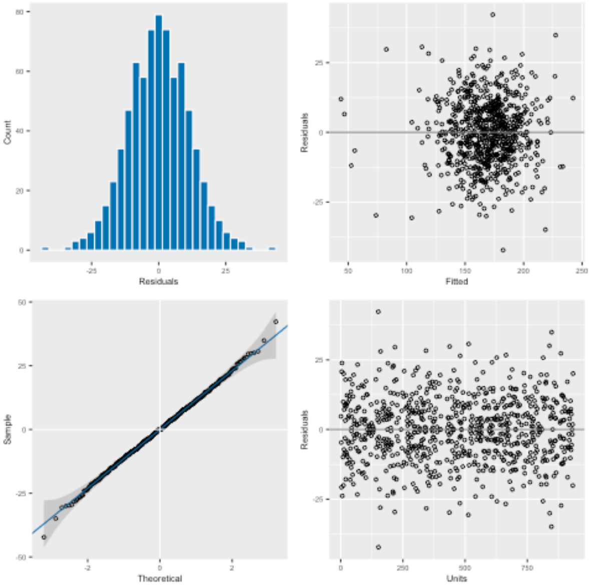

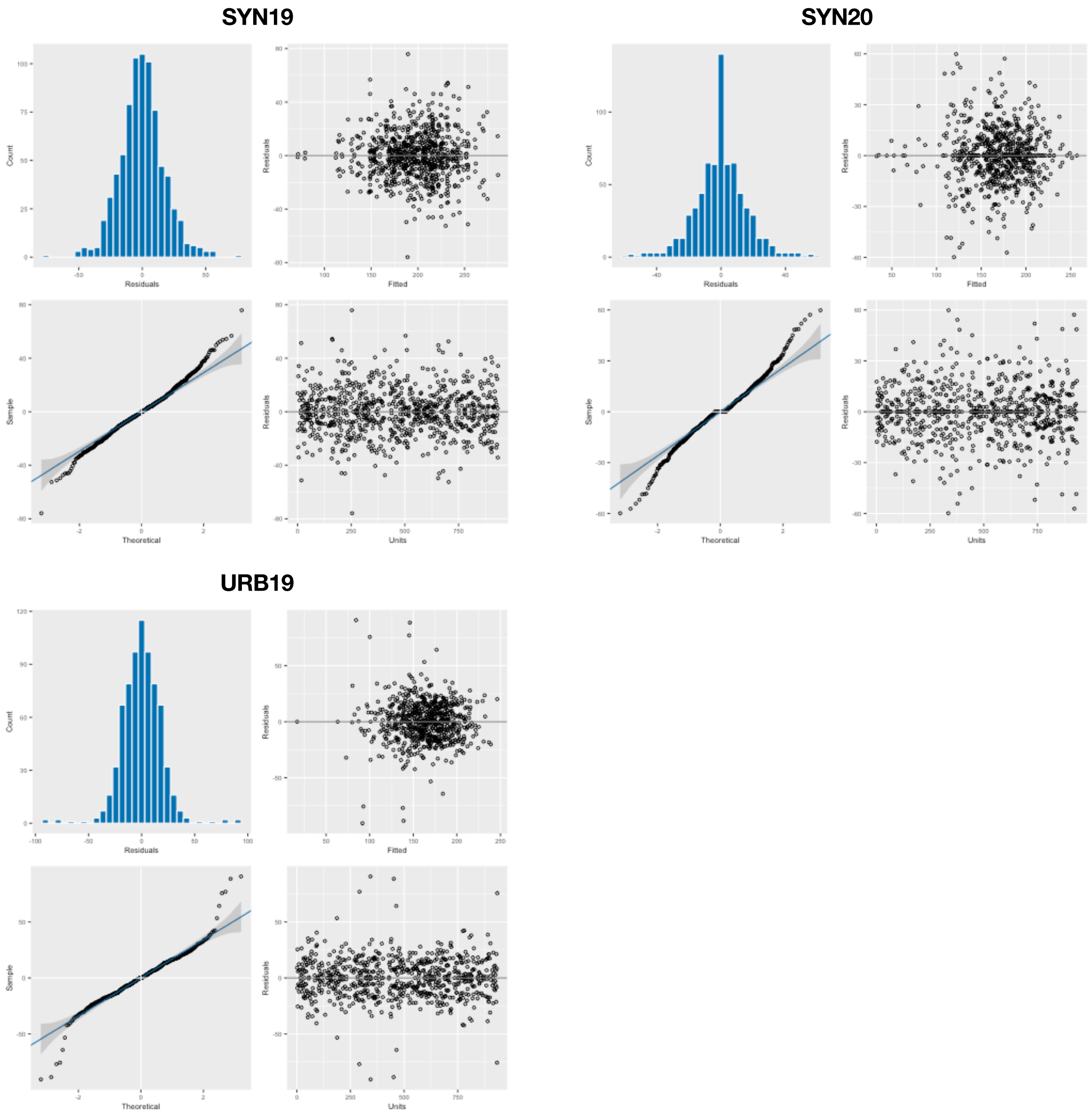


**Figure S1**. Residual distribution and quantile-quantile (Q-Q) plots for analysis of variance model used to generate best linear unbiased estimates within each environment.

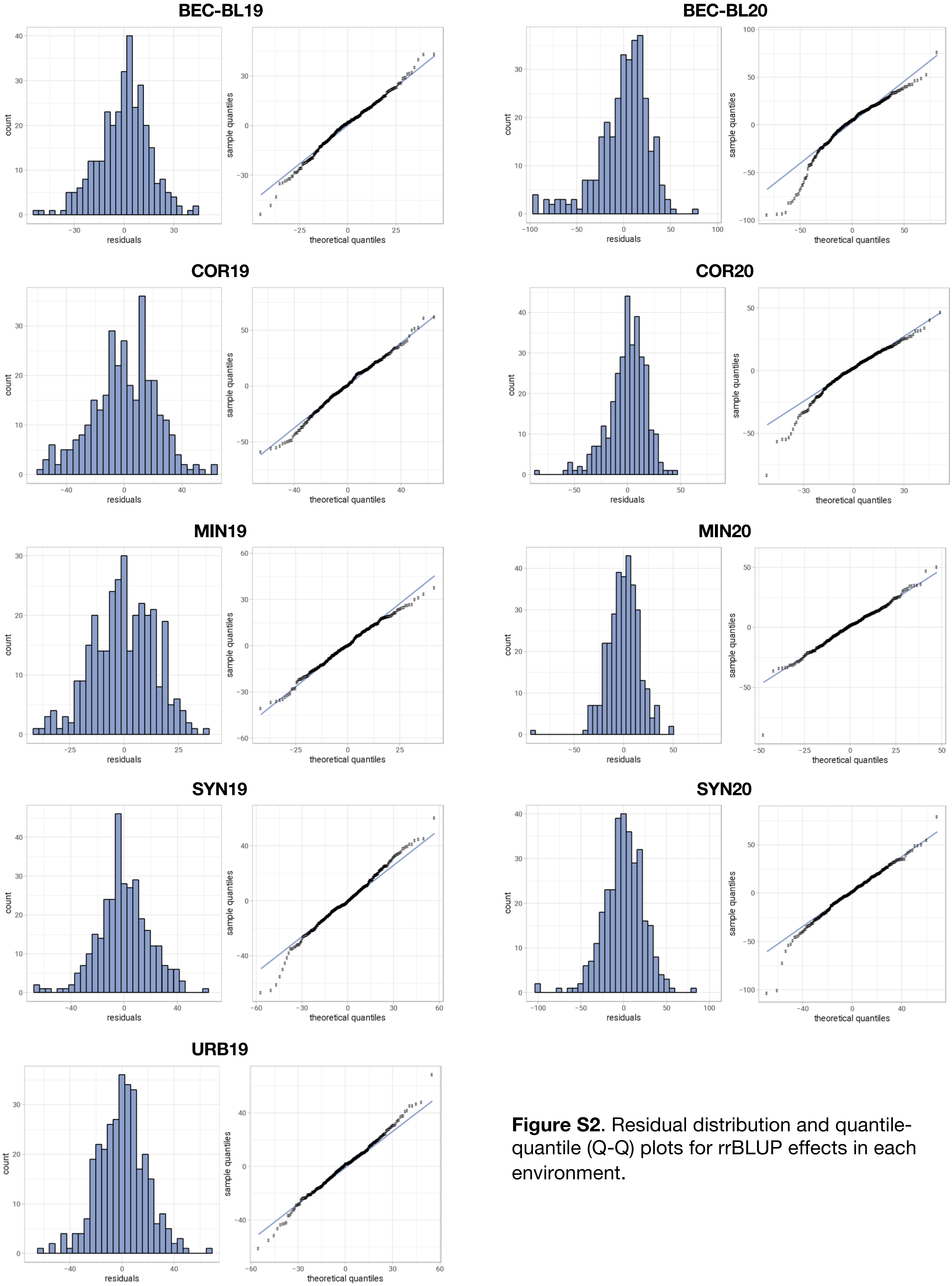


**Figure S2**. Residual distribution and quantile-quantile (Q-Q) plots for rrBLUP effects in each environment.

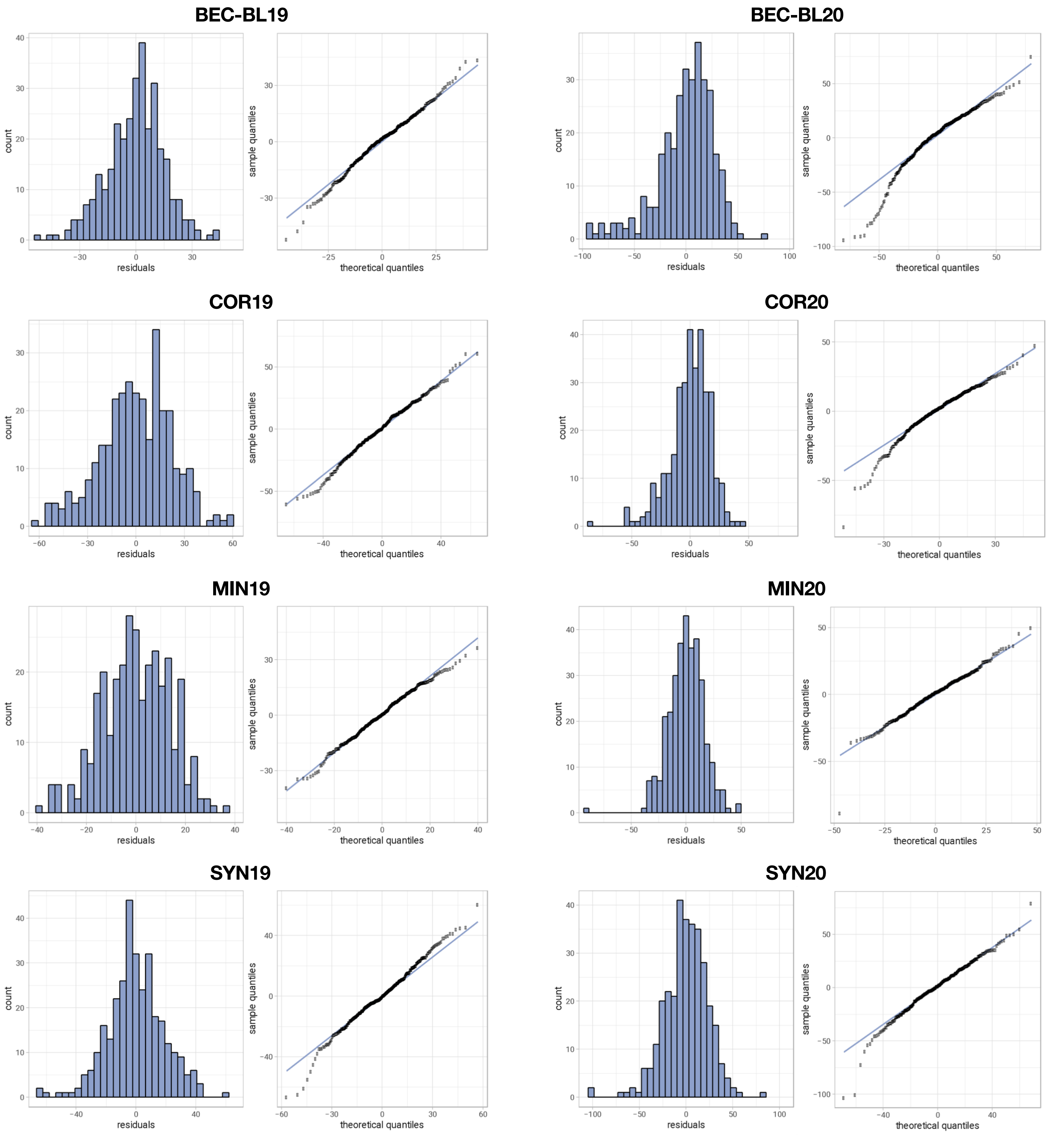


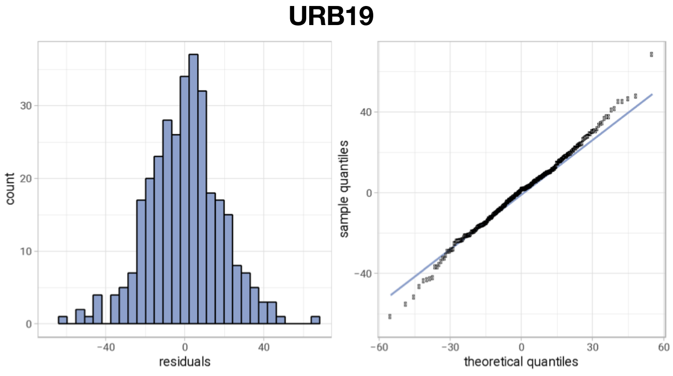


**Figure S3**. Residual distribution and quantile-quantile (Q-Q) plots for genome-wide association study (GWAS) effects in each environment.

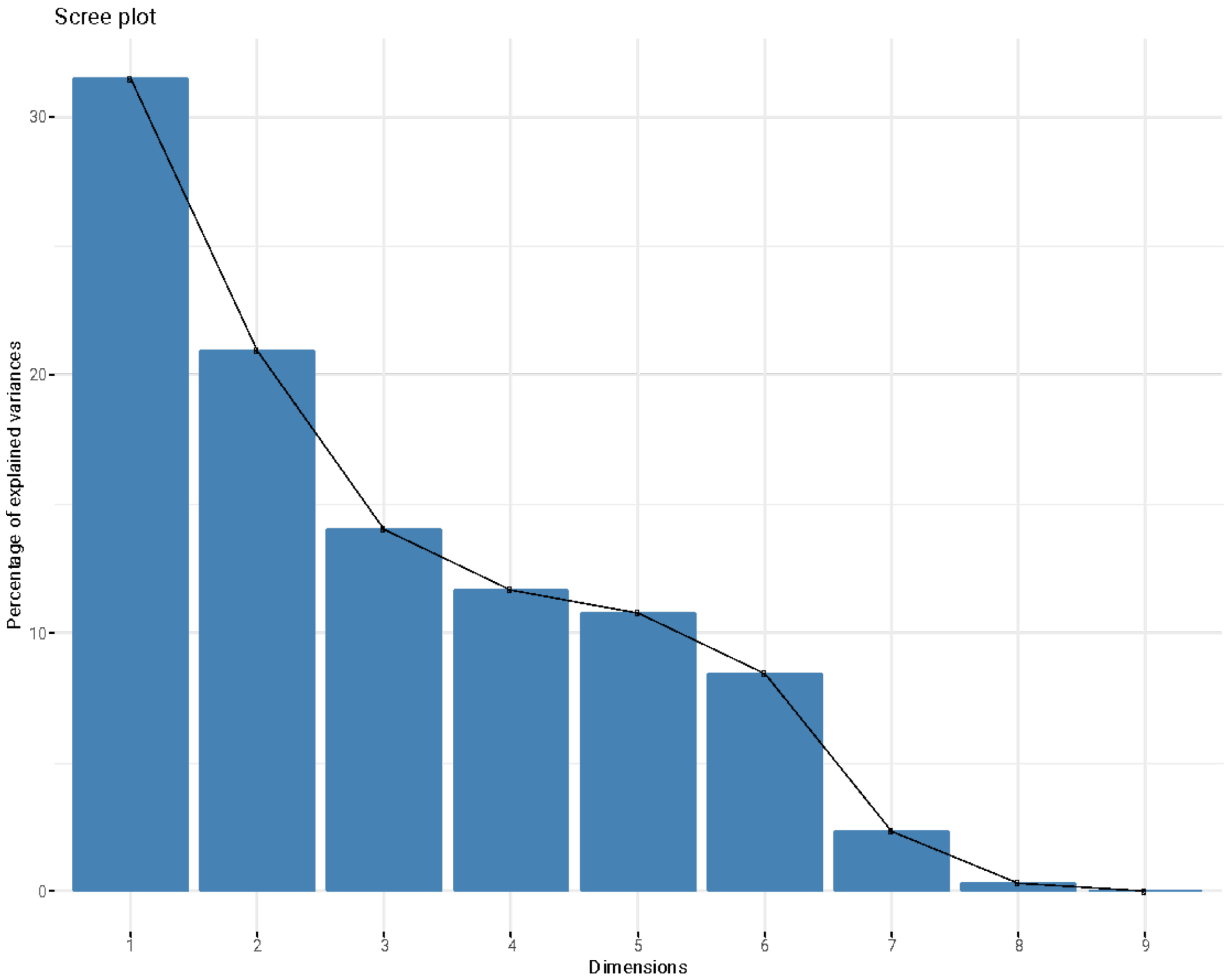


**Figure S4**. Percent variance explained by each principal component (PC) from 867 environmental parameters for for PCs 1-9.

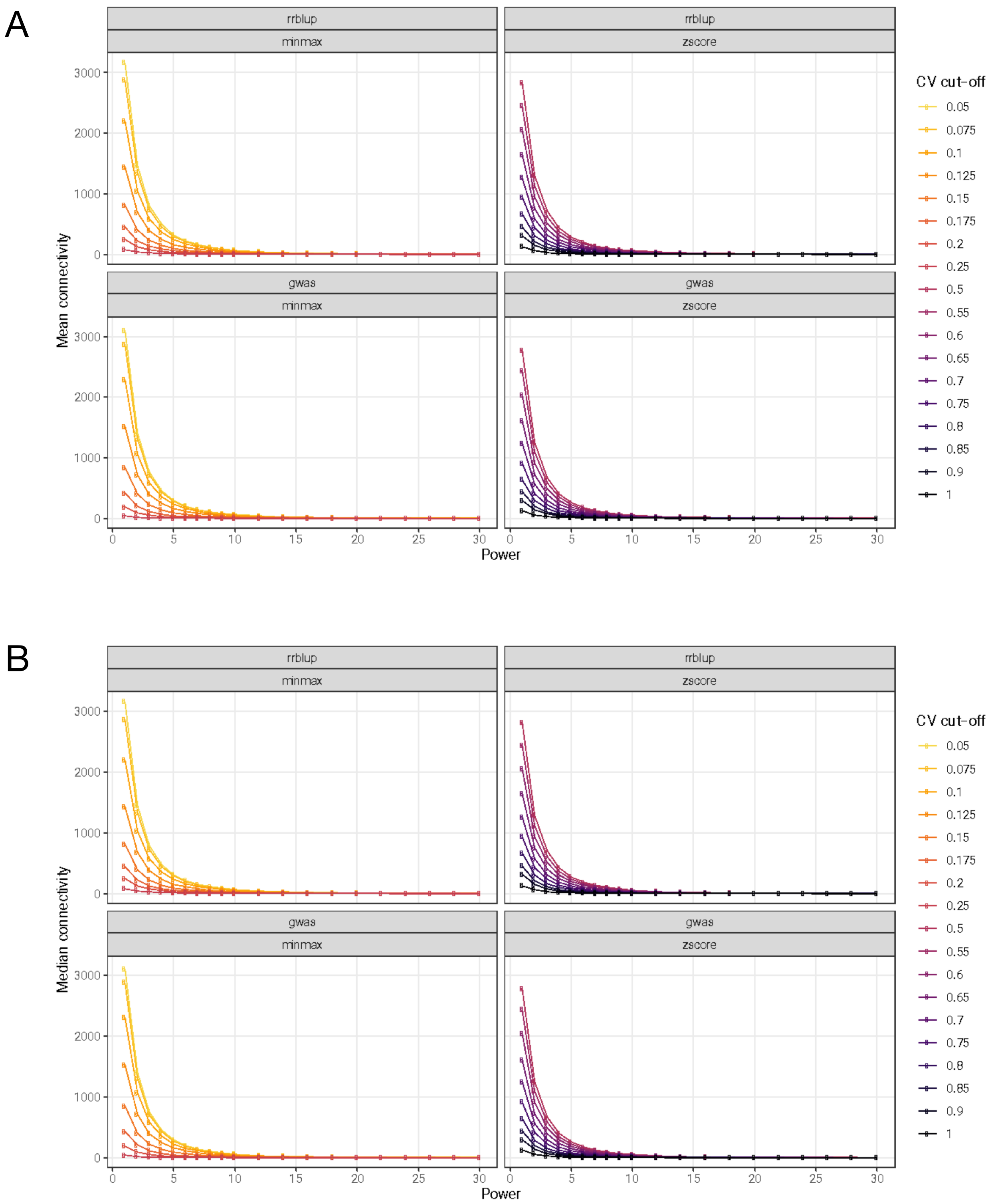


**Figure S5**. Relationship of marker effect estimation method, data normalization method, weighted gene co-expression network analysis (WGCNA) pipeline soft thresholding power level, and coefficient of variation cut-off with mean (A) and median (B) network connectivity score.

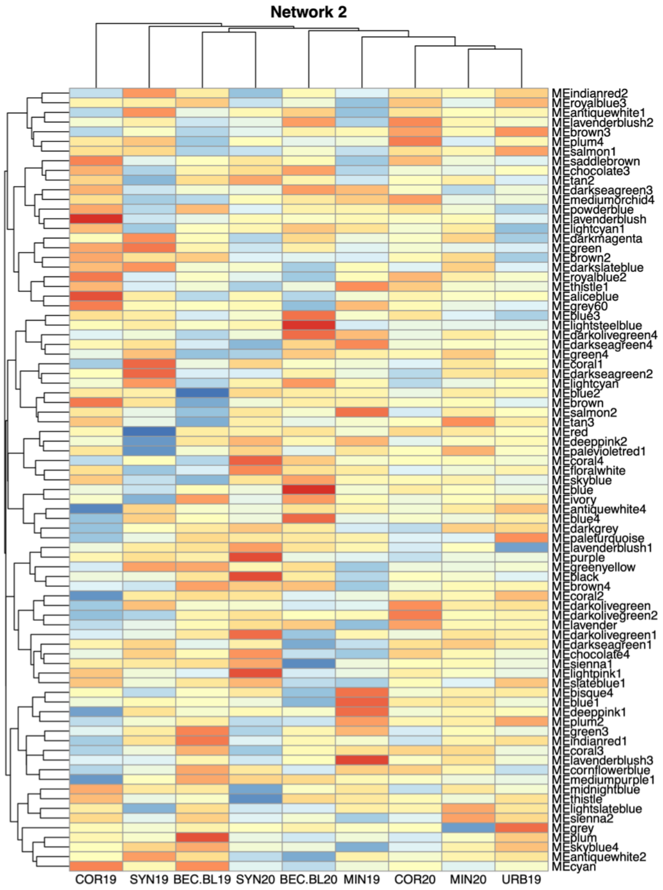
Network 2
0.5
0
−0.5
MEindianred2
MEroyalblue3
MEantiquewhite1
MElavenderblush2
MEbrown3
MEplum4
MEsalmon1
MEsaddlebrown
MEchocolate3
MEtan2
MEdarkseagreen3
MEmediumorchid4
MEpowderblue
MElavenderblush
MElightcyan1
MEdarkmagenta
MEgreen
MEbrown2
MEdarkslateblue
MEroyalblue2
MEthistle1
MEaliceblue
MEgrey60
MEblue3
MElightsteelblue
MEdarkolivegreen4
MEdarkseagreen4
MEgreen4
MEcoral1
MEdarkseagreen2
MElightcyan
MEblue2
MEbrown
MEsalmon2
MEtan3
MEred
MEdeeppink2
MEpalevioletred1
MEcoral4
MEfloralwhite
MEskyblue
MEblue
MEivory
MEantiquewhite4
MEblue4
MEdarkgrey
MEpaleturquoise
MElavenderblush1
MEpurple
MEgreenyellow
MEblack
MEbrown4
MEcoral2
MEdarkolivegreen
MEdarkolivegreen2
MElavender
MEdarkolivegreen1
MEdarkseagreen1
MEchocolate4
MEsienna1
MElightpink1
MEslateblue1
MEbisque4
MEblue1
MEdeeppink1
MEplum2
MEgreen3
MEindianred1
MEcoral3
MElavenderblush3
MEcornflowerblue
MEmediumpurple1
MEmidnightblue
MEthistle
MElightslateblue
MEsienna2
MEgrey
MEplum
MEskyblue4
MEantiquewhite2
MEcyan
COR19
SYN19
BEC.BL19
SYN20
BEC.BL20
MIN19
COR20
MIN20
URB19

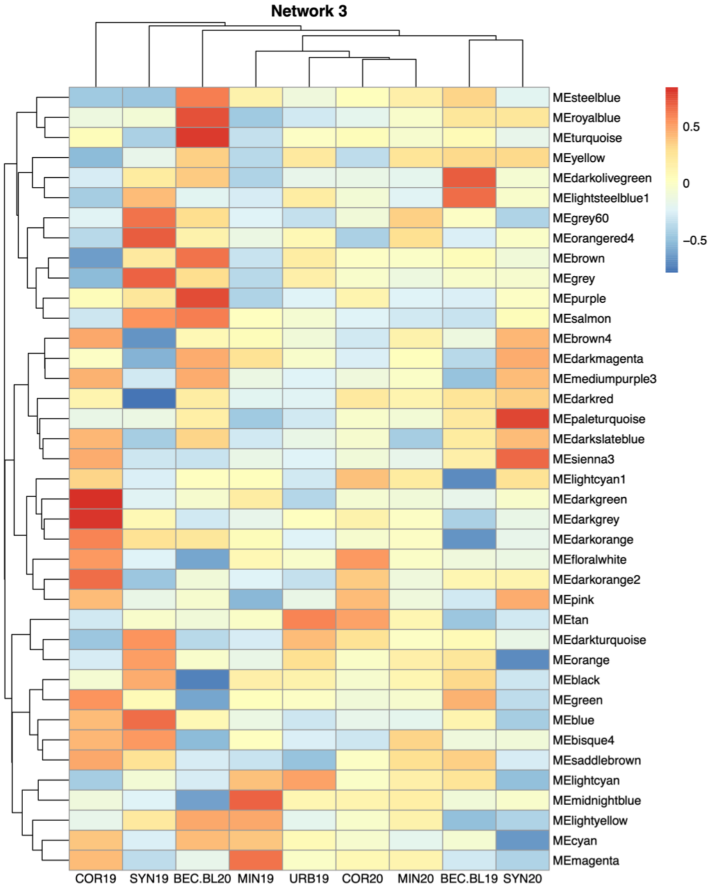
Network 3
0.5
0
−0.5
MEsteelblue
MEroyalblue
MEturquoise
MEyellow
MEdarkolivegreen
MElightsteelblue1
MEgrey60
MEorangered4
MEbrown
MEgrey
MEpurple
MEsalmon
MEbrown4
MEdarkmagenta
MEmediumpurple3
MEdarkred
MEpaleturquoise
MEdarkslateblue
MEsienna3
MElightcyan1
MEdarkgreen
MEdarkgrey
MEdarkorange
MEfloralwhite
MEdarkorange2
MEpink
MEtan
MEdarkturquoise
MEorange
MEblack
MEgreen
MEblue
MEbisque4
MEsaddlebrown
MElightcyan
MEmidnightblue
MElightyellow
MEcyan
MEmagenta
COR19
SYN19
BEC.BL20
MIN19
URB19
COR20
MIN20
BEC.BL19
SYN20

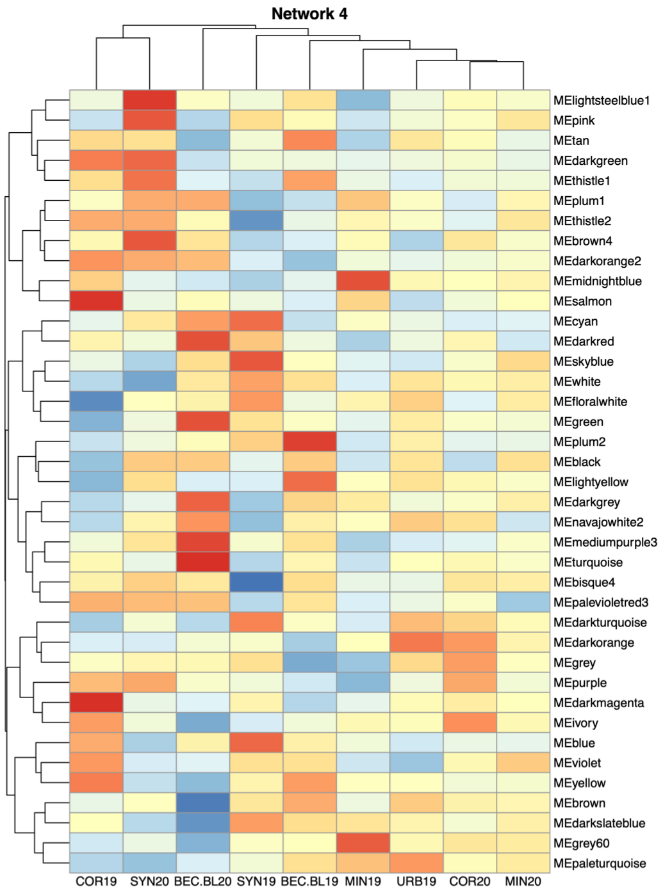
Network 4
0.5
0
−0.5
MElightsteelblue1
MEpink
MEtan
MEdarkgreen
MEthistle1
MEplum1
MEthistle2
MEbrown4
MEdarkorange2
MEmidnightblue
MEsalmon
MEcyan
MEdarkred
MEskyblue
MEwhite
MEfloralwhite
MEgreen
MEplum2
MEblack
MElightyellow
MEdarkgrey
MEnavajowhite2
MEmediumpurple3
MEturquoise
MEbisque4
MEpalevioletred3
MEdarkturquoise
MEdarkorange
MEgrey
MEpurple
MEdarkmagenta
MEivory
MEblue
MEviolet
MEyellow
MEbrown
MEdarkslateblue
MEgrey60
MEpaleturquoise
COR19
SYN20
BEC.BL20
SYN19
BEC.BL19
MIN19
URB19
COR20
MIN20

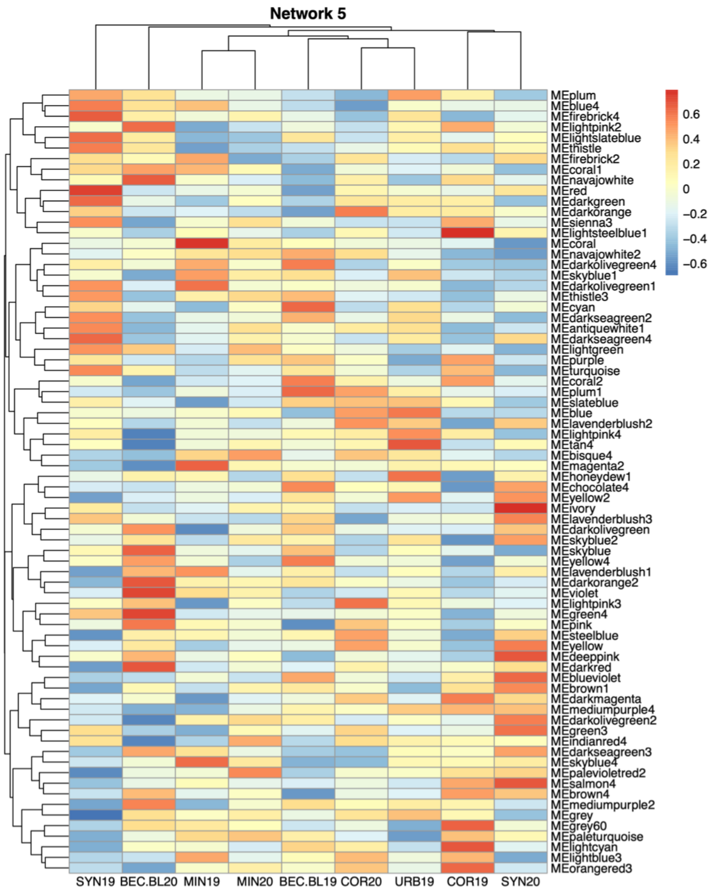
Network 5
0.6
0.4
0.2
0
−0.2
−0.4
−0.6
MEplum
MEblue4
MEfirebrick4
MElightpink2
MElightslateblue
MEthistle
MEfirebrick2
MEcoral1
MEnavajowhite
MEred
MEdarkgreen
MEdarkorange
MEsienna3
MElightsteelblue1
MEcoral
MEnavajowhite2
MEdarkolivegreen4
MEskyblue1
MEdarkolivegreen1
MEthistle3
MEcyan
MEdarkseagreen2
MEantiquewhite1
MEdarkseagreen4
MElightgreen
MEpurple
MEturquoise
MEcoral2
MEplum1
MEslateblue
MEblue
MElavenderblush2
MElightpink4
MEtan4
MEbisque4
MEmagenta2
MEhoneydew1
MEchocolate4
MEyellow2
MEivory
MElavenderblush3
MEdarkolivegreen
MEskyblue2
MEskyblue
MEyellow4
MElavenderblush1
MEdarkorange2
MEviolet
MElightpink3
MEgreen4
MEpink
MEsteelblue
MEyellow
MEdeeppink
MEdarkred
MEblueviolet
MEbrown1
MEdarkmagenta
MEmediumpurple4
MEdarkolivegreen2
MEgreen3
MEindianred4
MEdarkseagreen3
MEskyblue4
MEpalevioletred2
MEsalmon4
MEbrown4
MEmediumpurple2
MEgrey
MEgrey60
MEpaleturquoise
MElightcyan
MElightblue3
MEorangered3
SYN19
BEC.BL20
MIN19
MIN20
BEC.BL19
COR20
URB19
COR19
SYN20

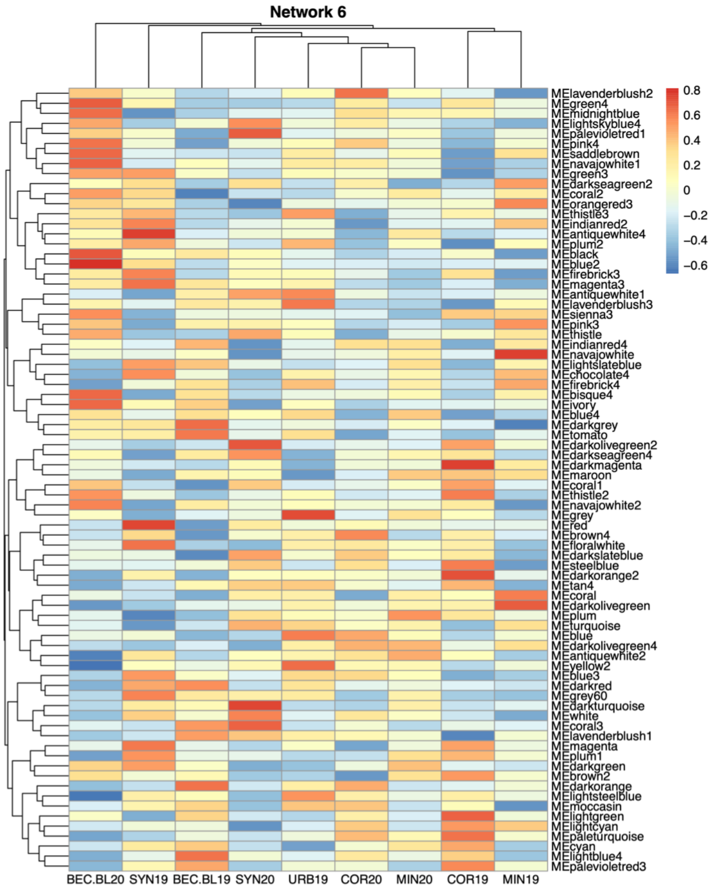
Network 6
0.8
0.6
0.4
0.2
0
−0.2
−0.4
−0.6
MElavenderblush2
MEgreen4
MEmidnightblue
MElightskyblue4
MEpalevioletred1
MEpink4
MEsaddlebrown
MEnavajowhite1
MEgreen3
MEdarkseagreen2
MEcoral2
MEorangered3
MEthistle3
MEindianred2
MEantiquewhite4
MEplum2
MEblack
MEblue2
MEfirebrick3
MEmagenta3
MEantiquewhite1
MElavenderblush3
MEsienna3
MEpink3
MEthistle
MEindianred4
MEnavajowhite
MElightslateblue
MEchocolate4
MEfirebrick4
MEbisque4
MEivory
MEblue4
MEdarkgrey
MEtomato
MEdarkolivegreen2
MEdarkseagreen4
MEdarkmagenta
MEmaroon
MEcoral1
MEthistle2
MEnavajowhite2
MEgrey
MEred
MEbrown4
MEfloralwhite
MEdarkslateblue
MEsteelblue
MEdarkorange2
MEtan4
MEcoral
MEdarkolivegreen
MEplum
MEturquoise
MEblue
MEdarkolivegreen4
MEantiquewhite2
MEyellow2
MEblue3
MEdarkred
MEgrey60
MEdarkturquoise
MEwhite
MEcoral3
MElavenderblush1
MEmagenta
MEplum1
MEdarkgreen
MEbrown2
MEdarkorange
MElightsteelblue
MEmoccasin
MElightgreen
MElightcyan
MEpaleturquoise
MEcyan
MElightblue4
MEpalevioletred3
BEC.BL20 SYN19 BEC.BL19 SYN20 URB19 COR20 MIN20 COR19 MIN19

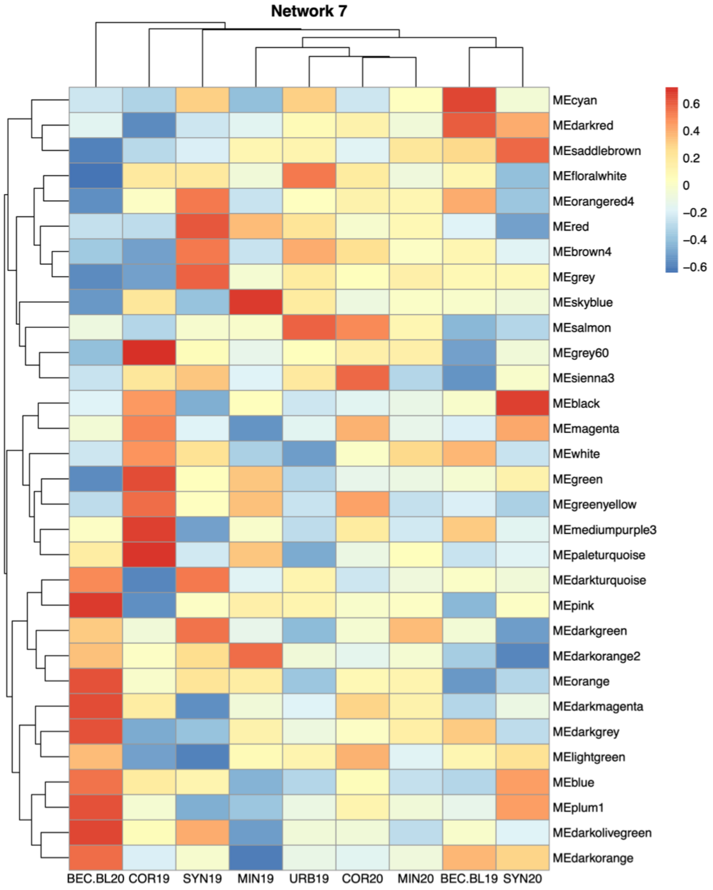
Network 7
0.6
0.4
0.2
0
−0.2
−0.4
−0.6
MEcyan
MEdarkred
MEsaddlebrown
MEfloralwhite
MEorangered4
MEred
MEbrown4
MEgrey
MEskyblue
MEsalmon
MEgrey60
MEsienna3
MEblack
MEmagenta
MEwhite
MEgreen
MEgreenyellow
MEmediumpurple3
MEpaleturquoise
MEdarkturquoise
MEpink
MEdarkgreen
MEdarkorange2
MEorange
MEdarkmagenta
MEdarkgrey
MElightgreen
MEblue
MEplum1
MEdarkolivegreen
MEdarkorange
BEC.BL20 COR19 SYN19 MIN19 URB19 COR20 MIN20 BEC.BL19 SYN20

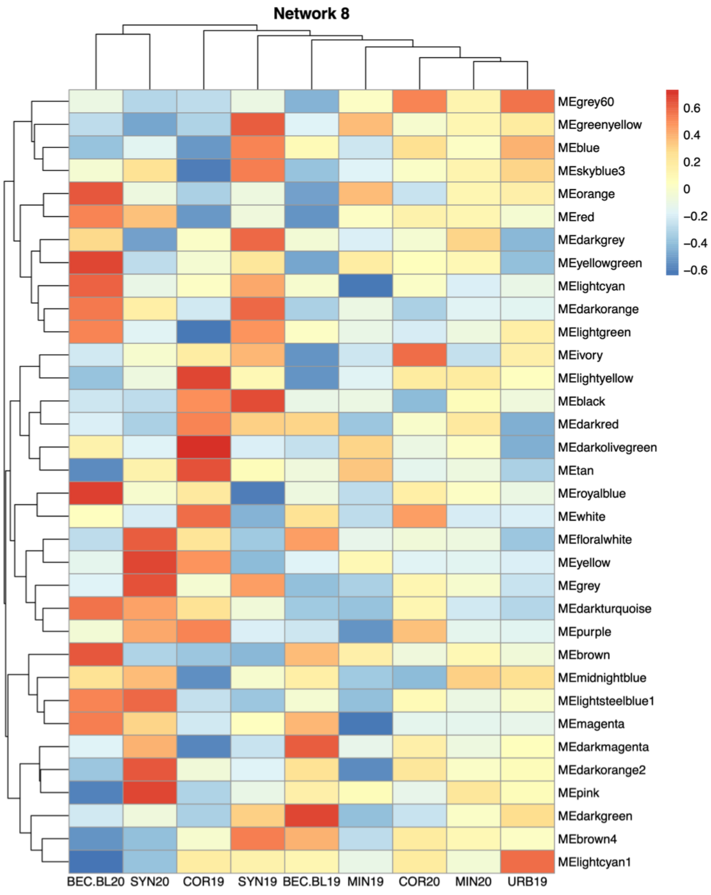
Network 8
0.6
0.4
0.2
0
−0.2
−0.4
−0.6
MEgrey60
MEgreenyellow
MEblue
MEskyblue3
MEorange
MEred
MEdarkgrey
MEyellowgreen
MElightcyan
MEdarkorange
MElightgreen
MEivory
MElightyellow
MEblack
MEdarkred
MEdarkolivegreen
MEtan
MEroyalblue
MEwhite
MEfloralwhite
MEyellow
MEgrey
MEdarkturquoise
MEpurple
MEbrown
MEmidnightblue
MElightsteelblue1
MEmagenta
MEdarkmagenta
MEdarkorange2
MEpink
MEdarkgreen
MEbrown4
MElightcyan1
BEC.BL20 SYN20 COR19 SYN19 BEC.BL19 MIN19 COR20 MIN20 URB19

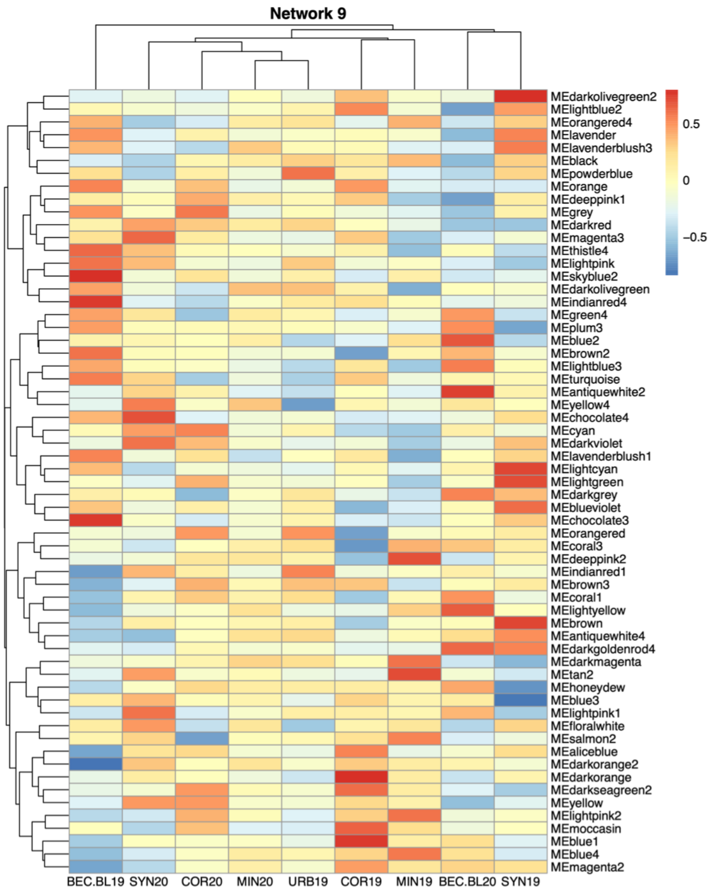
Network 9
0.5
0
−0.5
MEdarkolivegreen2
MElightblue2
MEorangered4
MElavender
MElavenderblush3
MEblack
MEpowderblue
MEorange
MEdeeppink1
MEgrey
MEdarkred
MEmagenta3
MEthistle4
MElightpink
MEskyblue2
MEdarkolivegreen
MEindianred4
MEgreen4
MEplum3
MEblue2
MEbrown2
MElightblue3
MEturquoise
MEantiquewhite2
MEyellow4
MEchocolate4
MEcyan
MEdarkviolet
MElavenderblush1
MElightcyan
MElightgreen
MEdarkgrey
MEblueviolet
MEchocolate3
MEorangered
MEcoral3
MEdeeppink2
MEindianred1
MEbrown3
MEcoral1
MElightyellow
MEbrown
MEantiquewhite4
MEdarkgoldenrod4
MEdarkmagenta
MEtan2
MEhoneydew
MEblue3
MElightpink1
MEfloralwhite
MEsalmon2
MEaliceblue
MEdarkorange2
MEdarkorange
MEdarkseagreen2
MEyellow
MElightpink2
MEmoccasin
MEblue1
MEblue4
MEmagenta2
BEC.BL19 SYN20 COR20 MIN20 URB19 COR19 MIN19 BEC.BL20 SYN19

**Network 10**

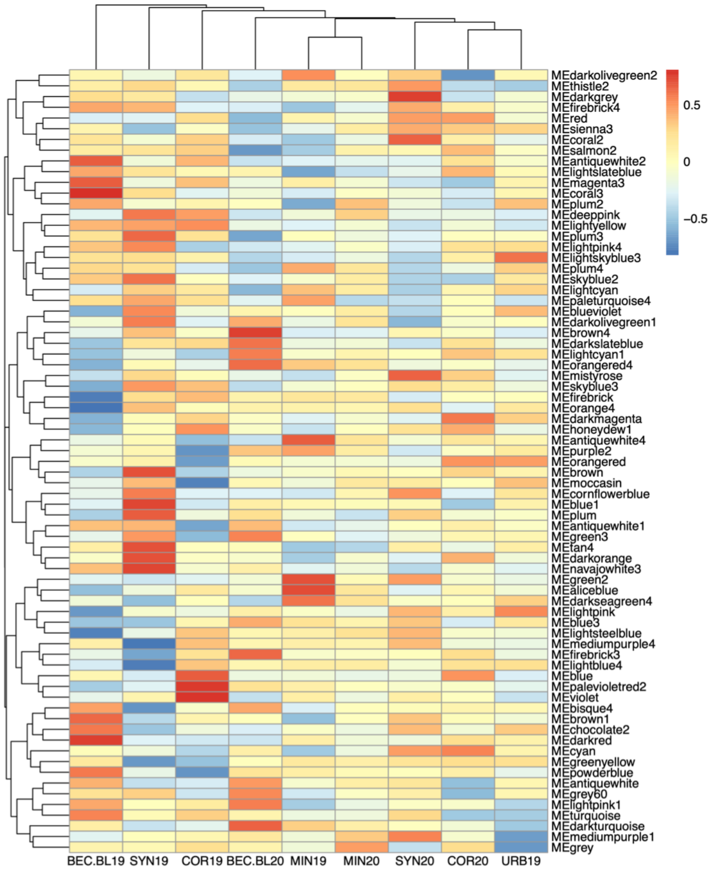


**Network 11**

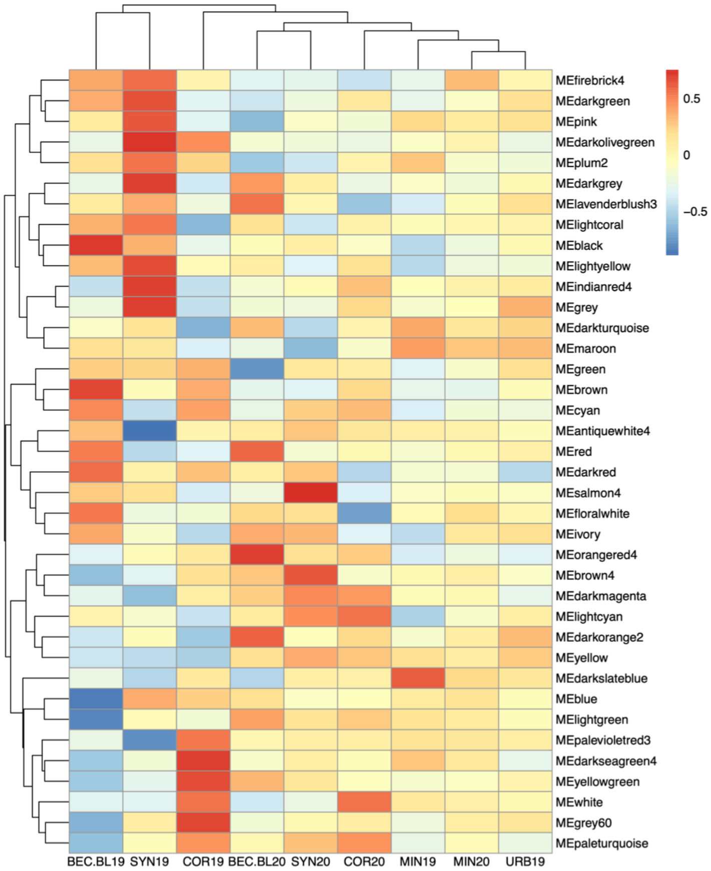


**Network 12**

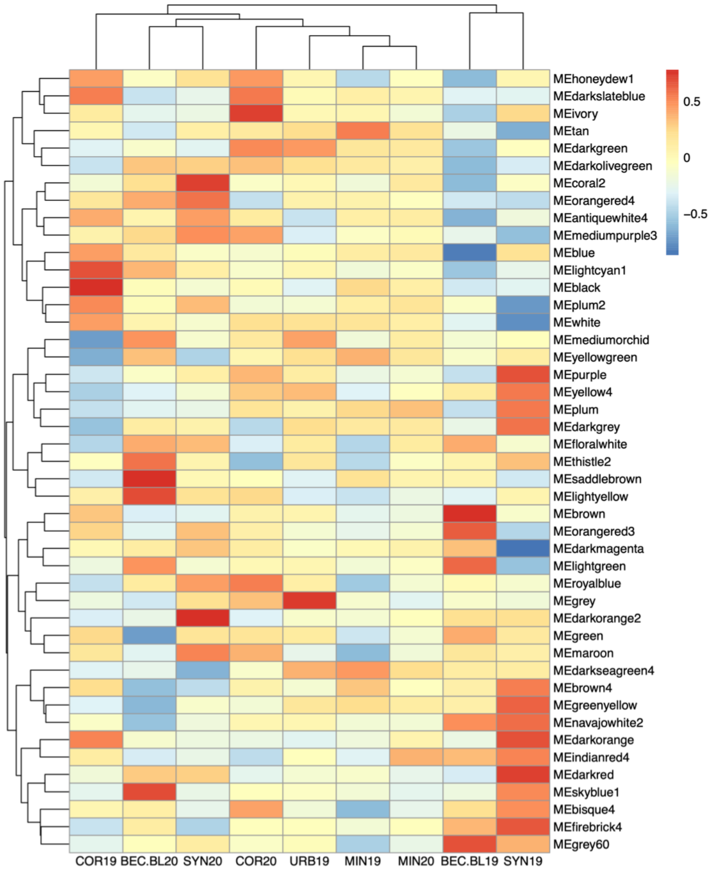


**Network 13**

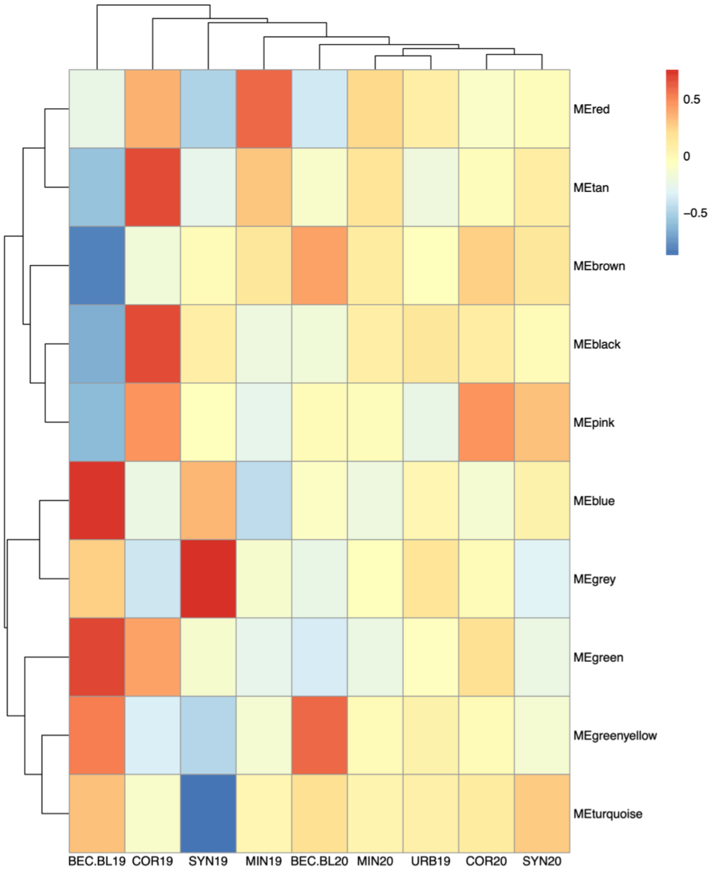

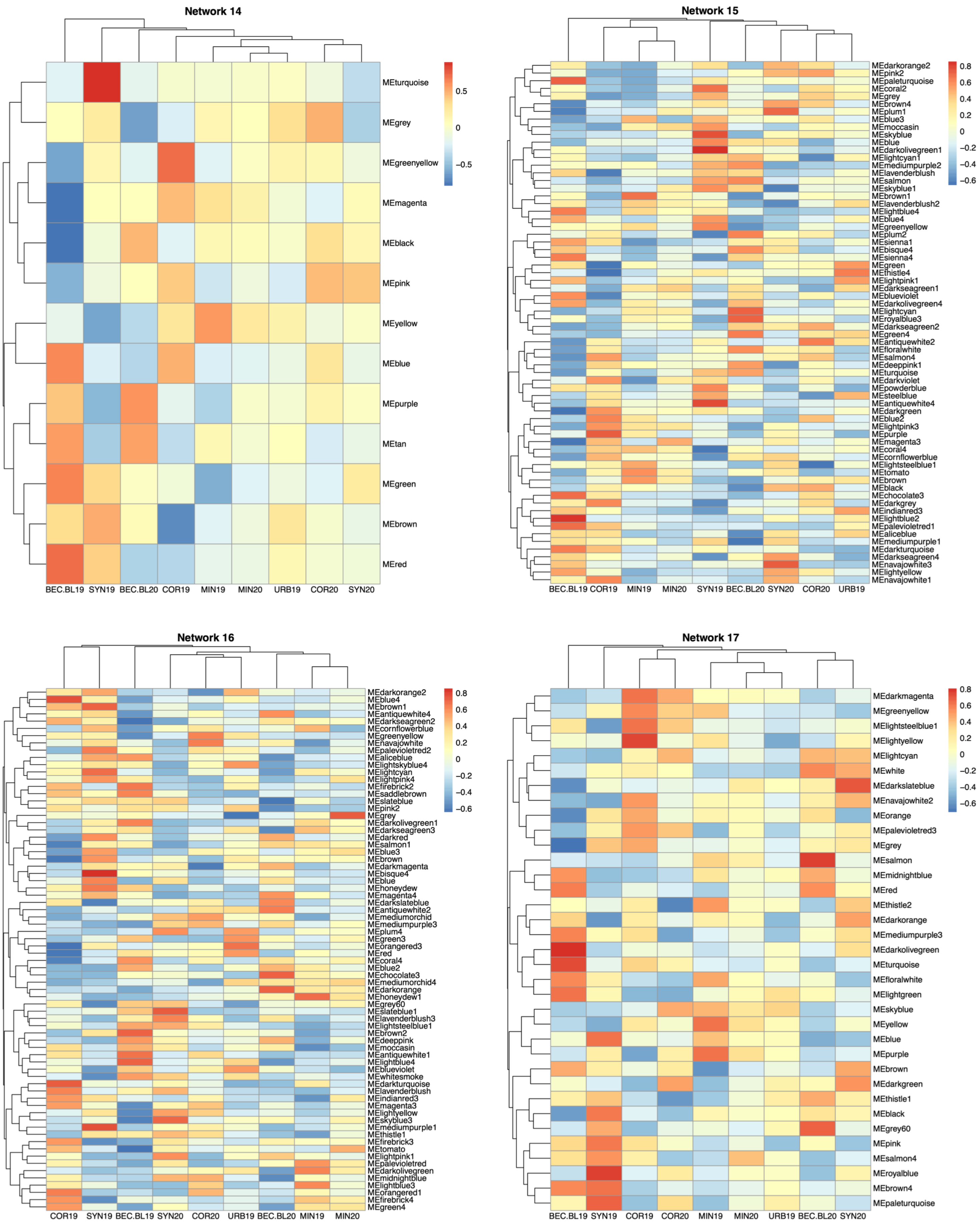
Network 14
MEturquoise
MEgrey
MEgreenyellow
MEmagenta
MEblack
MEpink
MEyellow
MEblue
MEpurple
MEtan
MEgreen
MEbrown
MEred
0.5
0
−0.5
BEC.BL19 SYN19 BEC.BL20 COR19 MIN19 MIN20 URB19 COR20 SYN20
Network 15
MEdarkorange2
MEpink2
MEpaleturquoise
MEcoral2
MEgrey
MEbrown4
MEplum1
MEblue3
MEmoccasin
MEskyblue
MEblue
MEdarkolivegreen1
MElightcyan1
MEmediumpurple2
MElavenderblush
MEsalmon
MEskyblue1
MEbrown1
MElavenderblush2
MElightblue4
MEblue4
MEgreenyellow
MEplum2
MEsienna1
MEbisque4
MEsienna4
MEgreen
MEthistle4
MElightpink1
MEdarkseagreen1
MEblueviolet
MEdarkolivegreen4
MElightcyan
MEroyalblue3
MEdarkseagreen2
MEgreen4
MEantiquewhite2
MEfloralwhite
MEsalmon4
MEdeeppink1
MEturquoise
MEdarkviolet
MEpowderblue
MEsteelblue
MEantiquewhite4
MEdarkgreen
MEblue2
MElightpink3
MEpurple
MEmagenta3
MEcoral4
MEcornflowerblue
MElightsteelblue1
MEtomato
MEbrown
MEblack
MEchocolate3
MEdarkgrey
MEindianred3
MElightblue2
MEpalevioletred1
MEaliceblue
MEmediumpurple1
MEdarkturquoise
MEdarkseagreen4
MEnavajowhite3
MElightyellow
MEnavajowhite1
0.8
0.6
0.4
0.2
0
−0.2
−0.4
−0.6
BEC.BL19 COR19 MIN19 MIN20 SYN19 BEC.BL20 SYN20 COR20 URB19
Network 16
MEdarkorange2
MEblue4
MEbrown1
MEantiquewhite4
MEdarkseagreen2
MEcornflowerblue
MEgreenyellow
MEnavajowhite
MEpalevioletred2
MEaliceblue
MElightskyblue4
MElightcyan
MElightpink4
MEfirebrick2
MEsaddlebrown
MEslateblue
MEpink2
MEgrey
MEdarkolivegreen1
MEdarkseagreen3
MEdarkred
MEsalmon1
MEblue3
MEbrown
MEdarkmagenta
MEbisque4
MEblue
MEhoneydew
MEmagenta4
MEdarkslateblue
MEantiquewhite2
MEmediumorchid
MEmediumpurple3
MEplum4
MEgreen3
MEorangered3
MEred
MEcoral4
MEblue2
MEchocolate3
MEmediumorchid4
MEdarkorange
MEhoneydew1
MEgrey60
MEslateblue1
MElavenderblush3
MElightsteelblue1
MEbrown2
MEdeeppink
MEmoccasin
MEantiquewhite1
MElightblue4
MEblueviolet
MEwhitesmoke
MEdarkturquoise
MElavenderblush
MEindianred3
MEmagenta3
MElightyellow
MEskyblue3
MEmediumpurple1
MEthistle1
MEfirebrick3
MEtomato
MElightpink1
MEpalevioletred
MEdarkolivegreen
MEmidnightblue
MElightblue3
MEorangered1
MEfirebrick4
MEgreen4
0.8
0.6
0.4
0.2
0
−0.2
−0.4
−0.6
COR19 SYN19 BEC.BL19 SYN20 COR20 URB19 BEC.BL20 MIN19 MIN20
Network 17
MEdarkmagenta
MEgreenyellow
MElightsteelblue1
MElightyellow
MElightcyan
MEwhite
MEdarkslateblue
MEnavajowhite2
MEorange
MEpalevioletred3
MEgrey
MEsalmon
MEmidnightblue
MEred
MEthistle2
MEdarkorange
MEmediumpurple3
MEdarkolivegreen
MEturquoise
MEfloralwhite
MElightgreen
MEskyblue
MEyellow
MEblue
MEpurple
MEbrown
MEdarkgreen
MEthistle1
MEblack
MEgrey60
MEpink
MEsalmon4
MEroyalblue
MEbrown4
MEpaleturquoise
0.8
0.6
0.4
0.2
0
−0.2
−0.4
−0.6
BEC.BL19 SYN19 COR19 COR20 MIN19 MIN20 URB19 BEC.BL20 SYN20

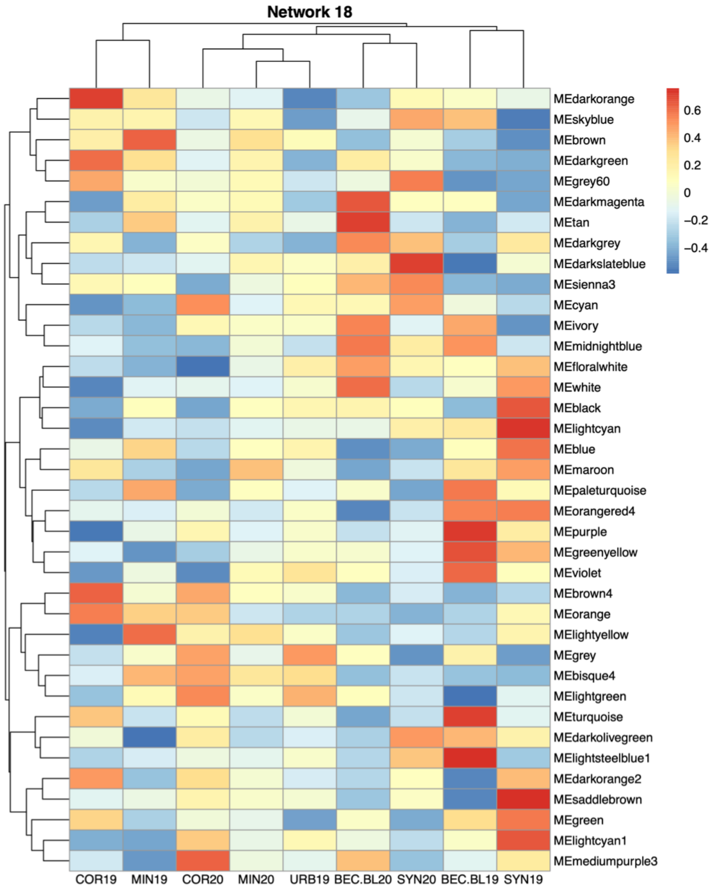


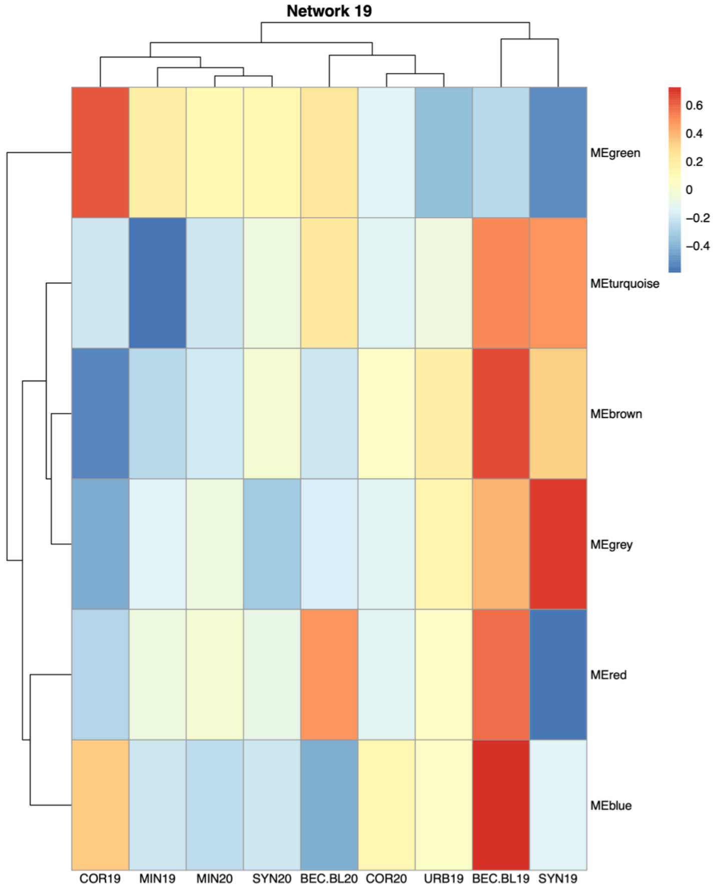


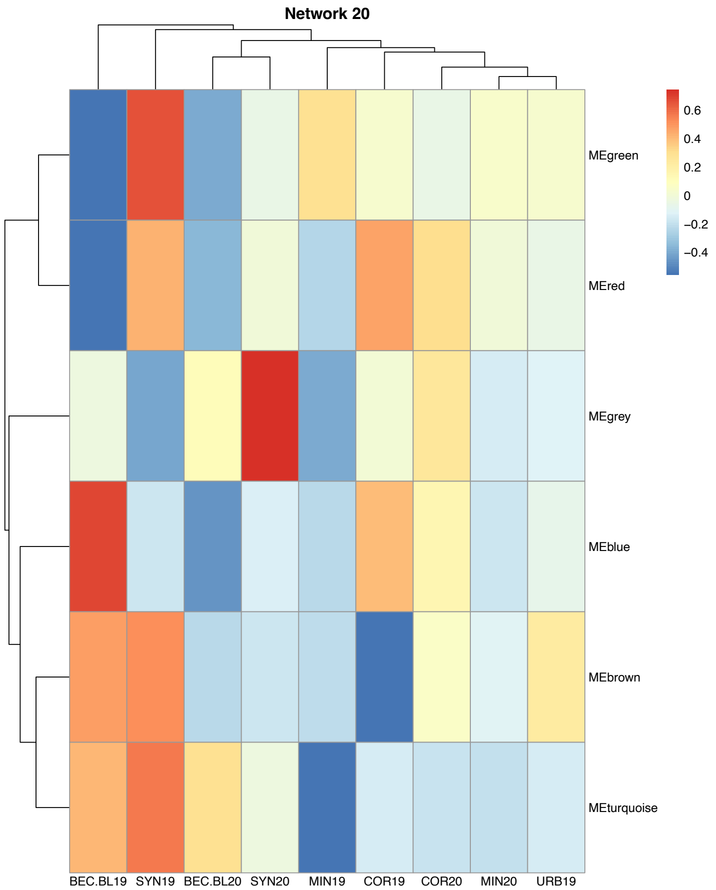


**Figure S6**. Heatmap of module eigenvalues (rows) across environments (columns) for networks 2-20. Higher eigenvalues are represented in red, while lower eigenvalues are in blue.

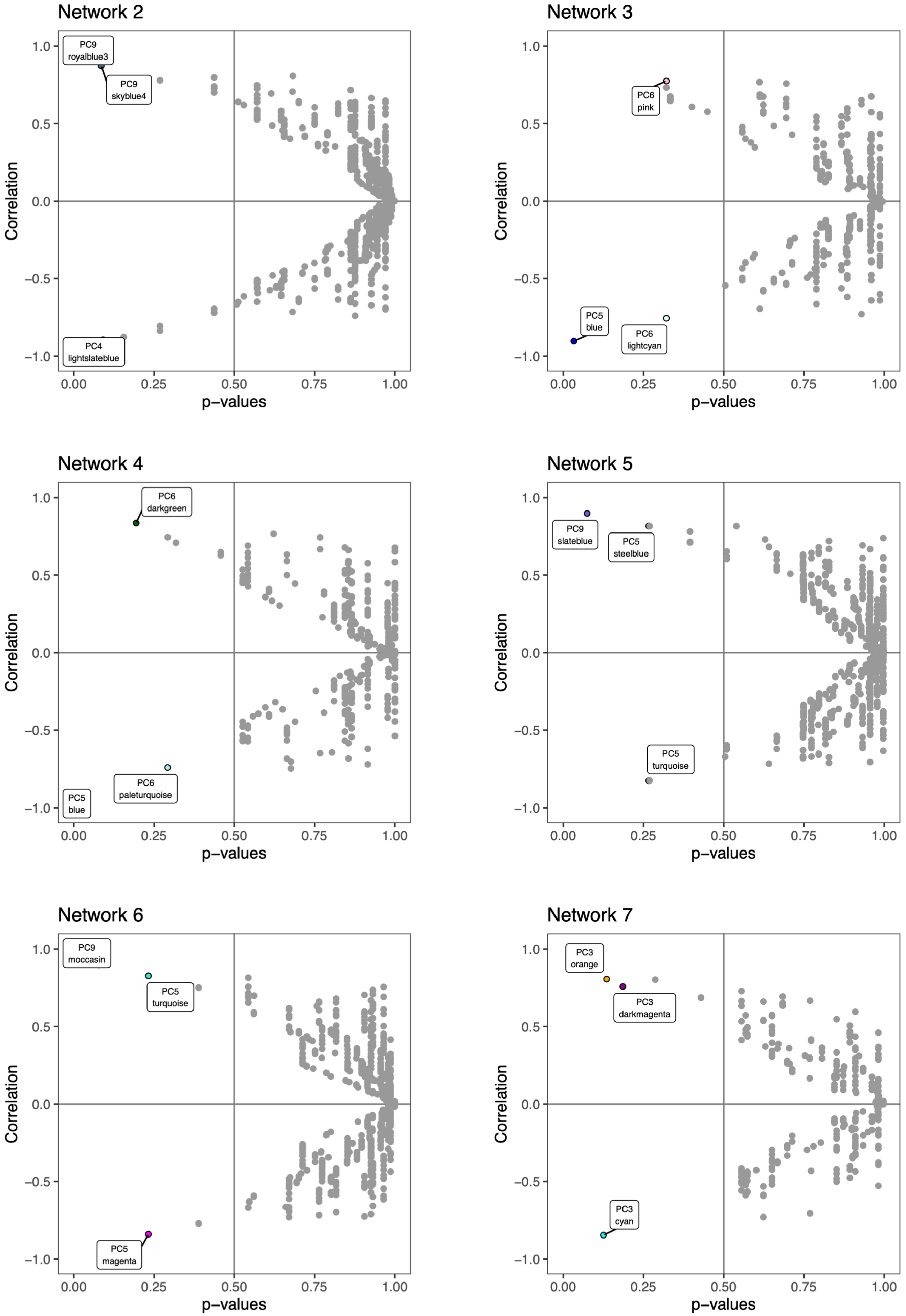
Network 2
PC9
royalblue3
PC9
skyblue4
PC4
lightslateblue
Network 3
PC6
pink
PC5
blue
PC6
lightcyan
Network 4
PC6
darkgreen
PC6
paleturquoise
PC5
blue
Network 5
PC9
slateblue
PC5
steelblue
PC5
turquoise
Network 6
PC9
moccasin
PC5
turquoise
PC5
magenta
Network 7
PC3
orange
PC3
darkmagenta
PC3
cyan
Correlation
p−values
1.0
0.5
0.0
−0.5
−1.0
0.00
0.25
0.50
0.75
1.00

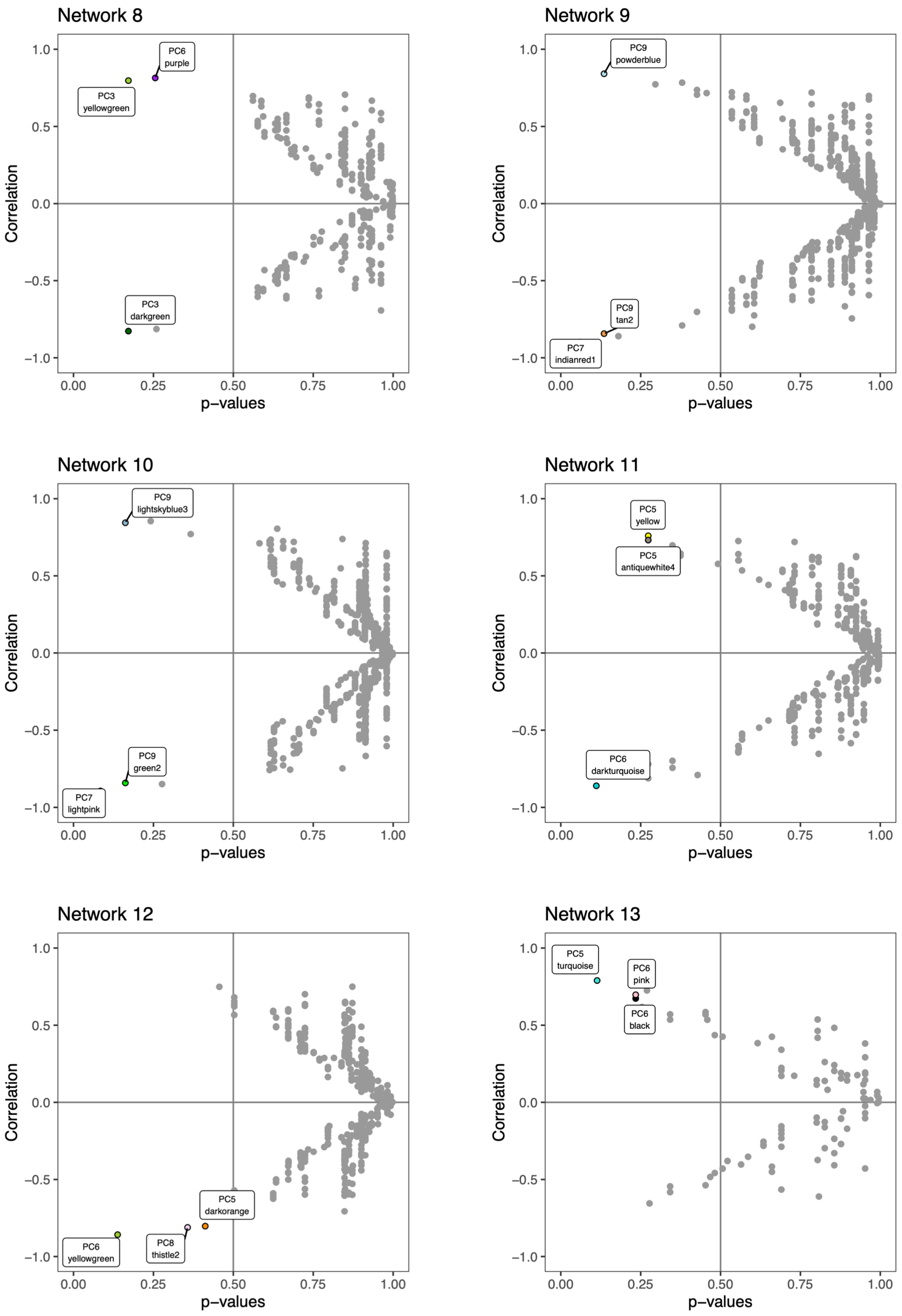
Network 8
PC6 purple
PC3 yellowgreen
PC3 darkgreen
Network 9
PC9 powderblue
PC9 tan2
PC7 indianred1
Network 10
PC9 lightskyblue3
PC9 green2
PC7 lightpink
Network 11
PC5 yellow
PC5 antiquewhite4
PC6 darkturquoise
Network 12
PC5 darkorange
PC8 thistle2
PC6 yellowgreen
Network 13
PC5 turquoise
PC6 pink
PC6 black
Correlation
p−values
1.0
0.5
0.0
−0.5
−1.0
0.00
0.25
0.50
0.75
1.00

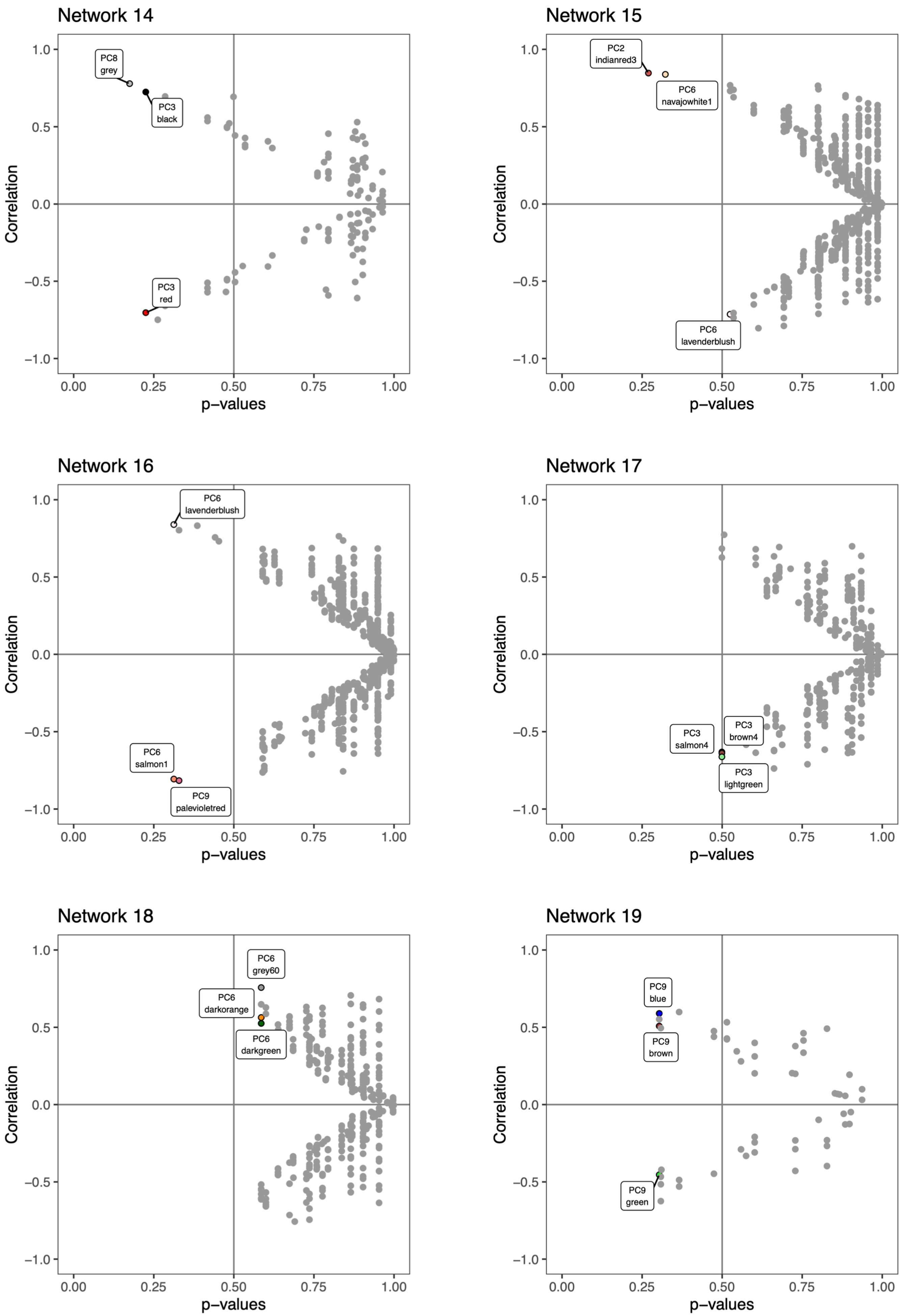

Network 14
PC8
grey
PC3
black
PC3
red
Network 15
PC2
indianred3
PC6
navajowhite1
PC6
lavenderblush
Network 16
PC6
lavenderblush
PC6
salmon1
PC9
palevioletred
Network 17
PC3
salmon4
PC3
brown4
PC3
lightgreen
Network 18
PC6
grey60
PC6
darkorange
PC6
darkgreen
Network 19
PC9
blue
PC9
brown
PC9
green
Correlation
p−values
1.0
0.5
0.0
−0.5
−1.0
0.00
0.25
0.50
0.75
1.00

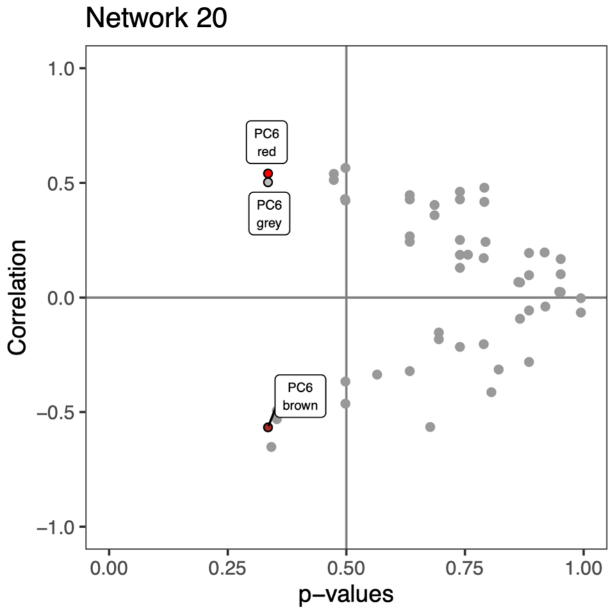


**Figure S7**. Correlation strength and p-value between the eigenvalues of each network module and PC (represented by circles) for networks 2-20.
